# A model of photosynthetic CO_2_ assimilation in C_3_ leaves accounting for respiration and energy recycling by the plastidial oxidative pentose phosphate pathway

**DOI:** 10.1101/2021.07.30.454461

**Authors:** Thomas Wieloch, Angela Augusti, Jürgen Schleucher

**Author notes:** Twitter: https://twitter.com/WielochThomas. ResearchGate: https://www.researchgate.net/profile/Thomas-Wieloch. GoogleScholar: https://scholar.google.de/citations?user=g0ZB7JMAAAAJ&hl=de&oi=sra. **Author, Email address** Thomas Wieloch Angela Augusti Jürgen Schleucher.

## Abstract

- Recently, we reported estimates of anaplerotic carbon flux through the oxidative pentose phosphate pathway (OPPP) in chloroplasts into the Calvin-Benson cycle. These estimates were based on intramolecular hydrogen isotope analysis of sunflower leaf starch. However, the isotope method is believed to underestimate actual flux at low atmospheric CO_2_ concentration (*C*_a_).
- Since the OPPP releases CO_2_ and reduces NADP^+^, it can be expected to affect leaf gas exchange under both rubisco- and RuBP-regeneration-limited conditions. Therefore, we expanded Farquhar-von Caemmerer-Berry models to account for OPPP metabolism. Based on model parameterisation with values from the literature, we estimated OPPP-related effects on leaf carbon and energy metabolism in the sunflowers analysed previously.
- We found that flux through the plastidial OPPP increases both above and below *C*_a_ ≈ 450 ppm (the condition the plants were acclimated to). This is qualitatively consistent with our previous isotope-based estimates, yet gas-exchange-based estimates are larger at low *C*_a_.
- We discuss our results in relation to regulatory properties of the plastidial and cytosolic OPPP, the proposed variability of CO_2_ mesophyll conductance, and the contribution of day respiration to the *A*/*C*_i_ curve drop at high *C*_a_. Furthermore, we critically examine the models and parameterisation and derive recommendations for follow-up studies.

## Introduction

Anaplerotic carbon flux into the Calvin-Benson cycle (CBC; for a list of abbreviations and symbols see Table 1) can occur by two metabolic pathways (Fig. 1). The first pathway (here denoted as cytosolic pathway) was proposed by Eicks *et al*. (2002) on discovering the xylulose 5-phosphate/phosphate translocator in *Arabidopsis thaliana*. This translocator enables exchange of pentose phosphates, and inorganic phosphate between chloroplasts and the cytosol. Screening *Arabidopsis* genome databases, Eicks *et al*. (2002) found genes encoding for the cytosolic oxidative branch of the pentose phosphate pathway (OPPP) as well as cytosolic ribulose-5-phosphate epimerase and ribose-5-phosphate isomerase. Genes encoding for cytosolic transketolase and transaldolase required to process pentose phosphates were missing. Because of the apparent lack of fate for pentose phosphates in the cytosol, Eicks *et al*. (2002) proposed their translocation into chloroplasts and injection into the CBC. Recently, studies on *Camelina sativa* leaf metabolism reported that this pathway carries high flux (≈ 5% relative to net CO_2_ assimilation, *A*) and supplies significant amounts of cytosolic NADPH (≈ 10% relative to *A*) at an atmospheric CO_2_ concentration (*C*_a_) of ≈ 400 ppm (Xu *et al*., 2022; Wieloch & Sharkey, 2022).

**Table 1.** List of abbreviations and symbols

| Abbreviation | Definition |
| --- | --- |
| CBC | Calvin-Benson cycle |
| F6P | fructose 6-phosphate |
| FvCB | Farquhar-von Caemmerer-Berry |
| G6P | glucose 6-phosphate |
| G6PD | glucose-6-phosphate dehydrogenase |
| OPPP | oxidative branch of the pentose phosphate pathway |
| PGI | phosphoglucose isomerase |
| Ru5P | ribulose 5-phosphate |
| RuBP | ribulose 1,5-bisphosphate |
| Symbol | Definition |
| $A$ | rate of net CO<sub>2</sub> assimilation |
| $A_c$ | rate of rubisco-limited CO<sub>2</sub> assimilation |
| $A_j$ | rate of RuBP-regeneration-limited CO<sub>2</sub> assimilation |
| $A_m$ | measured net CO<sub>2</sub> assimilation |
| $abs$ | absorptance of leaves |
| $C_a$ | atmospheric CO<sub>2</sub> concentration |
| $C_c$ | CO<sub>2</sub> partial pressure or concentration at the active site of rubisco |
| $C_i$ | CO<sub>2</sub> partial pressure or concentration in intercellular air spaces |
| $g_m$ | CO<sub>2</sub> mesophyll conductance |
| $I$ | incident photon flux |
| $I_2$ | incident quanta utilised in electron transport through photosystem II |
| $J$ | rate of linear electron transport |
| $J_{a,p}$ | rate of electron recycling by the plastidial anaplerotic pathway |
| $J_{max}$ | light-saturated rate of linear electron transport |
| $K_c$ | Michaelis-Menten constants of rubisco carboxylation |
| $K_o$ | Michaelis-Menten constants of rubisco oxygenation |
| $O_c$ | oxygen partial pressure at the active site of rubisco |
| $R_d$ | rate of day respiration |
| $R_{a,p}$ | rate of respiration by and flux through the plastidial anaplerotic pathway |
| $R_x$ | rate of respiration by pathways other than the plastidial anaplerotic pathway |
| $S_{c/o}$ | relative CO<sub>2</sub>/O<sub>2</sub> specificity of rubisco |
| $T$ | temperature |
| $V_c$ | rubisco carboxylation rate |
| $V_{cmax}$ | CO<sub>2</sub>-saturated maximum rate of rubisco carboxylation |
| $V_o$ | rubisco oxygenation rate |
| $\Gamma^*$ | CO<sub>2</sub>-compensation point in the absence of day respiration |
| $\Delta A_j$ | absolute effect of flux through the plastidial anaplerotic pathway on $A/C_i$ curves |
| $\theta$ | empirical curvature factor of the light response of electron transport |
| $\phi$ | $V_o/V_c$ ratio |

**Figure 1.**
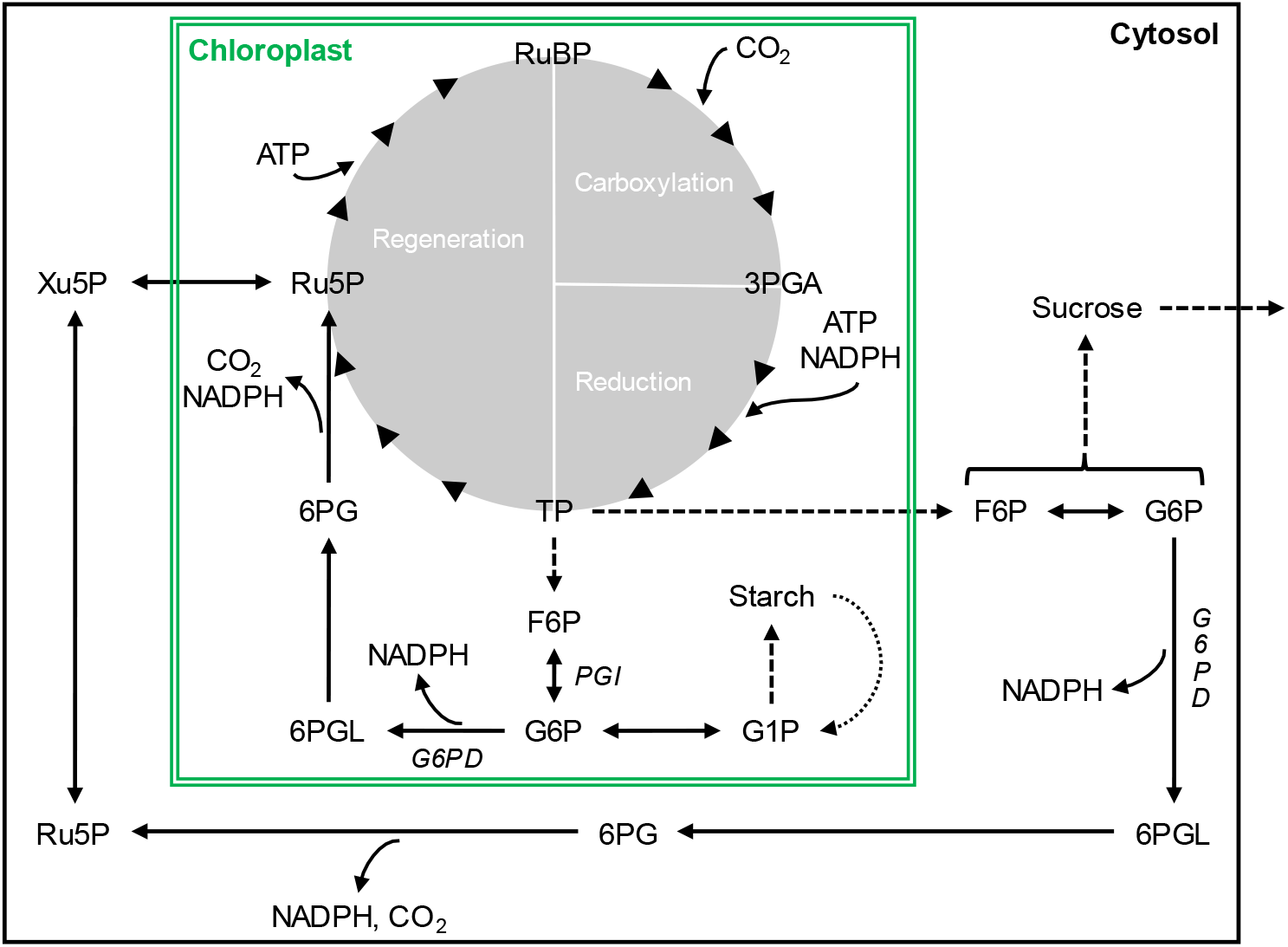
Anaplerotic carbon flux into the Calvin-Benson cycle. Dotted arrow, phosphorolytic starch breakdown. Abbreviations: 3PGA, 3-phosphoglycerate; 6PG, 6-phosphogluconate; 6PGL, 6-phosphogluconolactone; ATP, adenosine triphosphate; F6P, fructose 6-phosphate; FBP, fructose 1,6-bisphosphate; G1P, glucose 1-phosphate; G6P, glucose 6-phosphate; G6PD, glucose-6-phosphate dehydrogenase; NADPH, nicotinamide adenine dinucleotide phosphate; PGI, phosphoglucose isomerase; Ru5P, ribulose 5-phosphate; RuBP, ribulose 1,5-bisphosphate; TP, triose phosphate (glyceraldehyde 3-phosphate, dihydroxyacetone phosphate); Xu5P, Xylulose 5-phosphate. Chloroplast and cytosolic metabolism inside and outside the green box, respectively. Solid and dashed arrows represent transformations catalysed by a single enzyme or multiple enzymes, respectively.

The second pathway (here denoted as plastidial pathway) was proposed by Sharkey and Weise (2016). It involves glucose 6-phosphate (G6P) to ribulose 5-phosphate (Ru5P) conversion by the plastidial OPPP and injection of the latter into the CBC. For *Pinus nigra*, we reported evidence consistent with flux through this pathway under drought (Wieloch *et al*., 2018, 2022b). In sunflower, significant flux occurs when plants raised at *C*_a_ ≈ 450 ppm are moved into low or high *C*_a_ (≈ 180, 280, 700, 1500 ppm) whereas flux at *C*_a_ ≈ 450 ppm was not significantly greater than zero (Wieloch *et al*., 2022a; Wieloch, 2022). By contrast, Xu *et al*. (2021) reported, in *Camelina sativa*, significant flux occurs at *C*_a_ ≈ 400 ppm corresponding to the conditions the plants were raised in. However, several assumptions of this study were criticised (Wieloch, 2021) and, after implementing amendments, the result was not confirmed (Xu *et al*., 2022).

In sunflower, we estimated flux through the plastidial anaplerotic pathway based on a hydrogen isotope signal in leaf starch (the term isotope signal denotes systematic variability in relative isotope abundance) (Wieloch *et al*., 2022a; Wieloch, 2022). Assuming 50% of the net assimilated carbon becomes leaf starch (Sharkey *et al*., 1985), anaplerotic flux proceeds at > 7%, > 5%, 0%, ≈ 2%, and ≈ 5% relative to *A* at *C*_a_ ≈ 180, 280, 450, 700, and 1500 ppm, respectively (Wieloch, 2022). However, this method is thought to strongly underestimate flux at 180 and 280 ppm because much of the hydrogen isotope signal can be expected to not arrive in starch under low-*C*_a_ conditions (Wieloch *et al*., 2022a). Therefore, the present study aims first and foremost at developing an alternative method for estimating flux through the plastidial anaplerotic pathway and secondly at exploring its capabilities.

For each molecule Ru5P synthesised anaplerotically, one molecule CO_2_ is released from metabolism while two molecules NADP^+^ are reduced (Fig. 1). Thus, we hypothesise anaplerotic flux affects leaf gas exchange and can be traced by gas exchange modelling. To test this, we expand Farquhar-von Caemmerer-Berry (FvCB) photosynthesis models by terms accounting for anaplerotic respiration and energy recycling. Using gas exchange data of sunflower, we estimate anaplerotic flux rates and associated effects on leaf carbon and energy metabolism. These data are unique in that they were collected during the same experiment and from the same plants which previously provided isotope evidence for flux through the plastidial anaplerotic pathway (Wieloch *et al*., 2022a; Wieloch, 2022). The results are discussed (*inter alia*) in relation to (i) previously reported flux estimates, (ii) regulatory properties of the plastidial and cytosolic anaplerotic pathway, (iii) the proposed variability of CO_2_ mesophyll conductance (*g*_m_) with *C*_a_ and during photosynthetic induction, and (iv) the contribution of day respiration to the often-seen *A*/*C*_i_ curve drop at high *C*_a_ (*C*_i_, CO_2_ concentration in intercellular air spaces). We close with a paragraph about weaknesses of the developed models and the chosen parameterisation and future research directions.

## Theory

### Photosynthesis models accounting for anaplerotic flux into the Calvin-Benson cycle

In FvCB photosynthesis models (Farquhar *et al*., 1980), net CO_2_ assimilation is given as

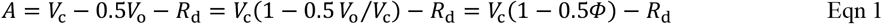

where *R*_d_ denotes day respiration, *V*_o_ and *V*_c_ denote the oxygenation and carboxylation rates of ribulose 1,5-bisphosphate (RuBP), respectively, and *Φ* denotes the *V*_o_/*V*_c_ ratio. Making day respiration by the plastidial anaplerotic pathway (*R*_a,p_) explicit yields

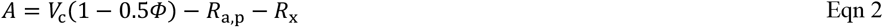

where *R*_x_ denotes day respiration by pathways other than the plastidial anaplerotic pathway. Please note that we also use *R*_a,p_ to denote flux through the plastidial anaplerotic pathway.

Different biochemical processes may exert control over *A*. First, rubisco-limited CO_2_ assimilation (*A*_c_) denotes assimilation limited by CO_2_ supply. Under these conditions, rubisco carboxylation rate is given as

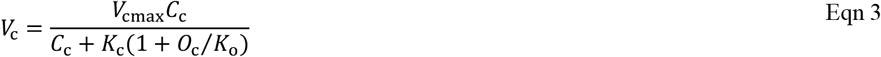

where *V*_cmax_ denotes the CO_2_-saturated maximum rate of rubisco carboxylation, *C*_c_ and *O*_c_ denote CO_2_ and O_2_ partial pressures at the active site of rubisco, respectively, and *K*_c_ and *K*_o_ denote the Michaelis-Menten constants of rubisco carboxylation and oxygenation, respectively (Farquhar *et al*., 1980). According to Farquhar and von Caemmerer (1982), *Φ* is related to the CO_2_-compensation point in the absence of day respiration (*Γ*\*) as

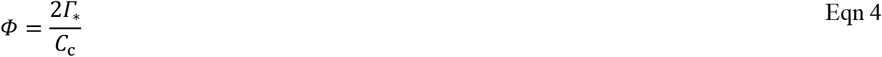

Using equations 3 and 4 to respectively remove *V*_c_ and *Φ* from equation 2 yields

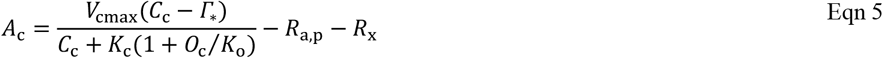

Second, RuBP-regeneration-limited CO_2_ assimilation (*A*_j_) presumably results from a shortage of incoming electrons supplied to the CBC and photorespiration as NADPH and ATP (Farquhar *et al*., 1980). Here, we assume NADPH rather than ATP supply is limiting. Electrons for NADP^+^ reduction come from the linear electron transport pathway (1 mol NADPH requires 2 mol electrons). Additionally, per 1 mol CO_2_ respired by anaplerotic pathways, 2 mol NADPH (4 mol electrons) are recycled (Fig. 1). However, only NADPH from the plastidial pathway can be recycled into the CBC and photorespiration. NADPH from the cytosolic pathway cannot enter chloroplasts (Wieloch & Sharkey, 2022). Thus, overall electron supply is given as

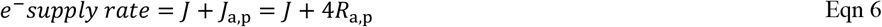

where *J* denotes the rate of linear electron transport, and *J*_a,p_ denotes the rate of electron recycling by the plastidial anaplerotic pathway. According to Farquhar *et al*. (1980), the rate of electron consumption by the CBC and photorespiration is given as

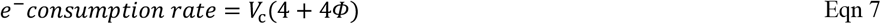

Balancing electron supply (Eqn. 6) with electron consumption (Eqn. 7) yields

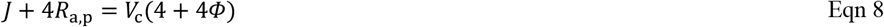

Solving equation 8 for *V*_c_ yields

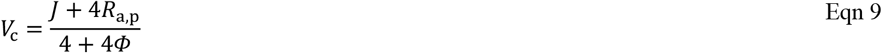

Using equation 9 to remove *V*_c_ from equation 2 yields

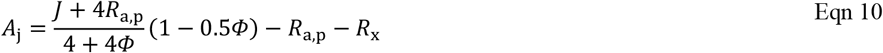

CO_2_ assimilation with electrons recovered by the plastidial anaplerotic pathway in the form of NADPH (Fig. 1) is given as

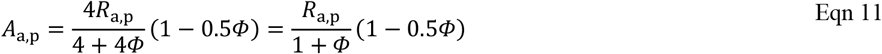

Using equation 4 to remove *Φ* from equation 10 yields

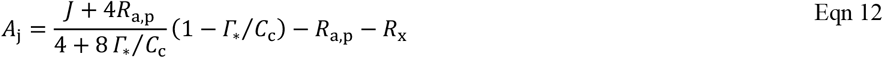

To quantify flux through the plastidial anaplerotic pathway and associated respiration, equation 12 is solved for *R*_a,p_ as

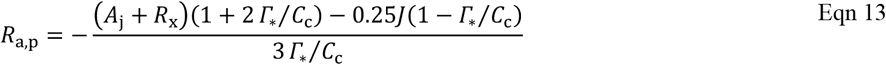

To calculate *V*_c_, *Φ* is removed from equation 9 using equation 4 as

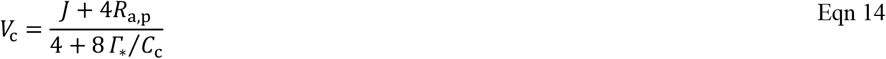

Using *Φ* (Eqn. 4), *V*_o_ is calculated as

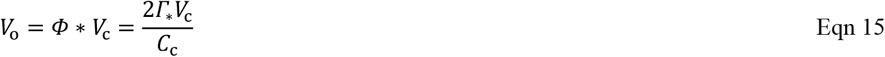

### Effects of respiration by the plastidial anaplerotic pathway on *A*/*C*_i_ curves

Respiration by the plastidial anaplerotic pathway can be expected to affect RuBP-regeneration-limited and rubisco-limited parts of *A*/*C*_i_ curves differently. In the following, effects associated with RuBP-regeneration-limited CO_2_ assimilation are discussed first. Anaplerotic flux of 1 mol is associated with the release of 1 mol CO_2_ and the supply of 2 mol NADPH (Fig. 1). In the complete absence of photorespiration, all this NADPH is used by the CBC (requires 2 NADPH per rubisco carboxylation) resulting in complete CO_2_ reassimilation. Thus, in this theoretical scenario, *A*_j_ does not change in response to *R*_a,p_, and gas exchange data do not convey any information on *R*_a,p_ no matter how large *R*_a,p_ is. By contrast, complete photorespiratory use of NADPH from the anaplerotic pathway (requires 2 NADPH per rubisco oxygenation) involves the release of 0.5 mol CO_2_ by photorespiration per 1 mol CO_2_ from the anaplerotic pathway (i.e., a total of 1.5 mol CO_2_ is released). Thus, in the RuBP-regeneration-limited case, *R*_a,p_ affects *A*/*C*_i_ curves because of photorespiration. The absolute effect of flux through the plastidial anaplerotic pathway on *A*/*C*_i_ curves (Δ*A*_j_) is given as

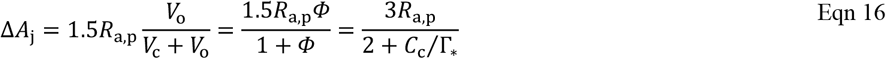

To calculate *R*_a,p_ from observed *A*_j_ offsets, equation 16 is solved for *R*_a,p_ as

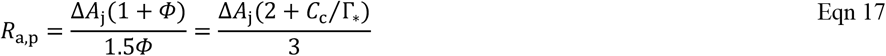

The relative effect of respiration by the plastidial anaplerotic pathway on *A*/*C*_i_ curves is given as

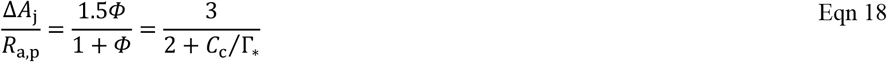

Hence, the larger *Φ*, the larger the Δ*A*_j_ / *R*_a,p_ ratio (Fig. 2, black line). That is, the higher photorespiration, the larger the effect of respiration by the plastidial anaplerotic pathway on *A*/*C*_i_ curves. At *Φ* = 2 (i.e., *C*_c_ = *Γ*\*, see Eqn. 4), Δ*A*_j_ = *R*_a,p_ (i.e., Δ*A*_j_ / *R*_a,p_ = 1). If *Φ* > 2, then Δ*A*_j_ > *R*_a,p_. If *Φ* < 2, then Δ*A*_j_ < *R*_a,p_.

**Figure 2.**
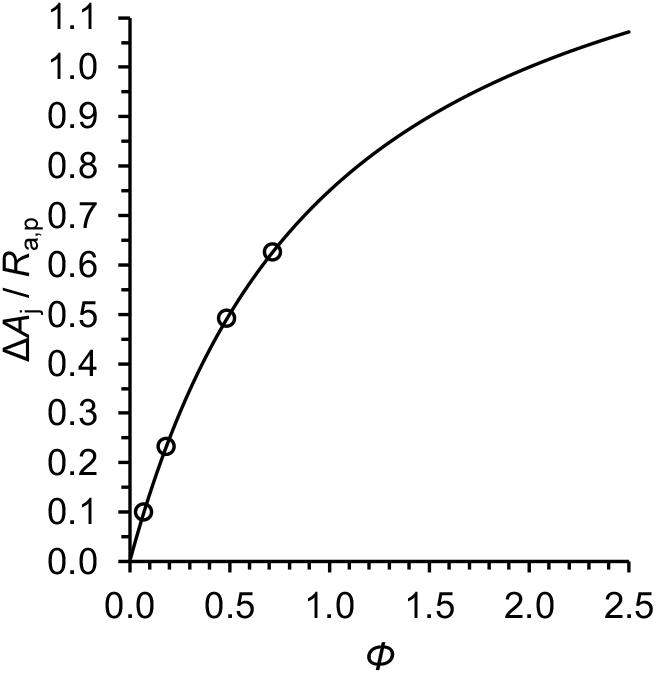
Relative effect of respiration by the plastidial anaplerotic pathway on *A*/*C*_i_ curves under RuBP-regeneration-limited conditions (Δ*A*_j_ / *R*_a,p_) as function of the oxygenation-to-carboxylation ratio of rubisco (*Φ*). Black line: Relationship based on theory (equation 18). Black circles: Δ*A*_j_ / *R*_a,p_ ratios pertaining to gas exchange data presented here where Δ*A*_j_ denotes the observed difference between modelled and measured RuBP-regeneration-limited CO_2_ assimilation (Fig. 4b, red line versus black dots, respiration by the plastidial anaplerotic pathway not considered by the model) and *R*_a,p_ denotes respiration by the plastidial anaplerotic pathway estimated based on equation 13.

Under rubisco-limited CO_2_ assimilation, RuBP concentrations are above saturation level. Hence, NADPH from the anaplerotic pathway can be expected to exert no control on *A*_c_. Shifts in *R*_a,p_ will cause *A*_c_ shifts of the same size.

### NADPH and ATP demand of plastidial carbon metabolism

NADPH demand by the CBC and photorespiration is given as (Farquhar *et al*., 1980)

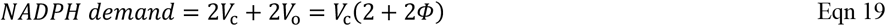

This demand is met by NADPH supply from the light reactions. In the presence of flux through the plastidial anaplerotic pathway, NADPH demand from the light reactions is reduced as

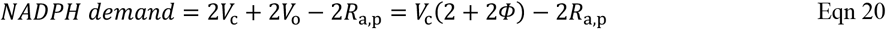

because anaplerotic flux of 1 mol is associated with the supply of 2 mol NADPH (Fig. 1).

Upon carboxylation of 1 mol RuBP, the CBC requires 3 mol ATP (Farquhar & von Caemmerer, 1982). Upon oxygenation of 1 mol RuBP, photorespiration requires 3 mol ATP in chloroplasts (1.5 mol by phosphoglycerate kinase, 1 mol by phosphoribulokinase, and 0.5 mol by glycerate kinase). This accounting assumes that potentially harmful ammonium released by the mitochondrial glycine decarboxylase complex is immediately recapture by mitochondrial (not plastidial) glutamine synthetase and then transported as glutamine to chloroplasts (Taira *et al*., 2004; Buchanan *et al*., 2015). ATP required by mitochondrial glutamine synthetase can be expected to come from mitochondrial oxidative phosphorylation. Hence, ATP demand by the CBC and photorespiration from the light reactions is given as

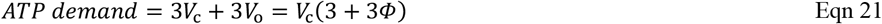

Combining equations 20 and 21, the ratio at which the CBC and photorespiration require ATP and NADPH is given as

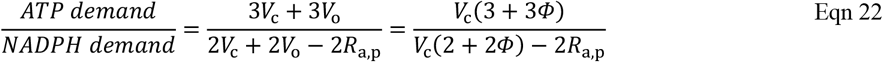

Evidently, the CBC and photorespiration have the same plastidial cofactor demands with a plastidial ATP-to-NADPH demand ratio of 1.5. The ratio is independent of *Φ*. However, NADPH demands from the light reactions vary with *R*_a,p_ resulting in varying ATP-to-NADPH demands from the light reactions.

## Materials and Methods

### Samples and leaf gas exchange measurements

The sunflower (*Helianthus annuus*) samples were described previously (Ehlers *et al*., 2015). In brief, the plants were raised in a greenhouse at *C*_a_ ≈ 450 ppm and a light intensity of 300-400 μmol photons m^−2^ s^−1^ (16 h photoperiod, 21% O_2_). After 7-8 weeks, they were transferred to a growth chamber (*T* = 22 °C : 18 °C, *rH* = 60% : 70%, day : night), kept in darkness for 1 day to drain the starch reserves (see Notes S3 in Wieloch *et al*., 2022a), and subsequently grown for 2 days in groups of 8 at *C*_a_ of either ≈ 180, 280, 450, 700, or 1500 ppm (300-400 μmol photons m^−2^ s^−1^, 16 h photoperiod).

Leaf gas exchange measurements were described previously (Ehlers *et al*., 2015; see Notes S1 in Wieloch *et al*., 2022a). In brief, the measurements were performed with a previously described system (Laisk & Edwards, 1997) under conditions similar to those in the growth chamber (*C*_a_ = 173, 269, 433, 675, 1431 ppm; *T* = 22 ± 0.5 °C; 300 μmol photons m^−2^ s^−1^, 21% O_2_). *C*_i_ values obtained from gas exchange measurements (136, 202, 304, 516, 1282 ppm) are similar to *C*_i_ values estimated for the 2-day growth chamber experiments (140, 206, 328, 531, and 1365 ppm) (see Notes S2 in Wieloch *et al*., 2022a). To constrain *V*_cmax_, we additionally measured gas exchange at 900 μmol photons m^−2^ s^−1^, *C*_a_ = 173, 269, 432, 674, 1430 ppm corresponding to *C*_i_ = 126, 188, 264, 450, 1204 ppm.

### Parameterisation of photosynthesis models

*C*_c_ was calculated according to Fick’s first law as

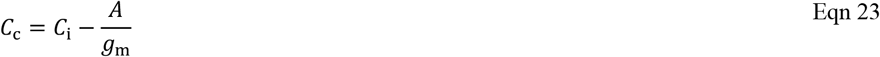

using *g*_m_ = 0.45 mol CO_2_ m^−2^ s^−1^ reported for sunflower at 20 °C, 600 μmol photons m^−2^ s^−1^ (Schäufele *et al*., 2011). For rubisco extracted from sunflower, Genkov *et al*. (2010) reported *K*_c_ = 19 μM CO_2_, *K*_o_ = 640 μM O_2_, and a relative CO_2_/O_2_ specificity (*S*_c/o_) of 77 [M/M] at 25 °C. *Γ*\* at 25 °C was calculated based on published procedures (von Caemmerer, 2000) as

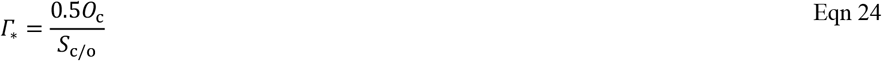

with *O*_c_ = 256 μM. To convert from concentration to partial pressure, solubilities for CO_2_ of 0.0334 mol l^−1^ bar^−1^ and O_2_ of 0.00126 mol l^−1^ bar^−1^ were used. Since correction factors for sunflower are unavailable, we used *Q*_10_ values reported in Table 2.3 of von Caemmerer (2000) and equation 2.32 in von Caemmerer (2000) to correct for leaf temperature (22 °C). This gave *K*_c_ = 447 μbar CO_2_, *K*_o_ = 439 mbar O_2_, and *Γ*\* = 45.3 μbar CO_2_. To estimate *V*_cmax_, equation 5 was fitted to gas exchange data obtained at 900 μmol photons m^−2^ s^−1^, *C*_a_ = 173, 269, 432 ppm (Fig. 3). Under standard conditions (400 ppm CO_2_, 21% O_2_, 20-25 °C), *R*_d_ usually proceeds at ≈ 5% relative to the rate of *A* (Tcherkez *et al*., 2017; Xu *et al*., 2022). Fixing *R*_a,p_ + *R*_x_ at this rate (= 0.83 μmol CO_2_ m^−2^ s^−1^), we found *V*_cmax_ = 83.2 μmol m^−2^ s^−1^. This is similar to a previously reported estimate for sunflower of 80 μmol m^−2^ s^−1^ at 20 °C (Jacob & Lawlor, 1991; Wullschleger, 1993).

**Table 2.** Rates of day respiration in sunflower leaves.

| $C_a$<br>[ppm] | $A_m$<br>[$\mu\text{mol m}^{-2} \text{s}^{-1}$] | $A_j - A_m$ <sup>(a)</sup><br>[$\mu\text{mol m}^{-2} \text{s}^{-1}$] | $R_{a,p}$ <sup>(b)</sup><br>[$\mu\text{mol m}^{-2} \text{s}^{-1}$] | $R_{a,p} / A_m$<br>% | $R_{a,p} / A_m$ <sup>(d)</sup><br>% | $R_x$ <sup>(e)</sup><br>[$\mu\text{mol m}^{-2} \text{s}^{-1}$] | $R_x / A_m$<br>% |
| --- | --- | --- | --- | --- | --- | --- | --- |
| $\approx 180$ | 5.13 | 1.73 | 2.75 | 53.7 | > 7 | 0.96 | 18.8 |
| $\approx 280$ | 8.46 | 1.20 | 2.44 | 28.9 | > 5 | 0.96 | 11.4 |
| $\approx 450$ | 12.29 | 0 | 0 | 0 | 0 | 0.96 | 7.8 |
| $\approx 700$ | 14.88 | 0.19 | 0.80 <sup>(c)</sup> | 5.3 <sup>(c)</sup> | $\approx 2$ | 1.08 | 7.2 |
| $\approx 1500$ | 17.33 | 0.51 | 5.03 <sup>(c)</sup> | 29.0 <sup>(c)</sup> | $\approx 5$ | 1.38 | 8.0 |

Plants were grown in a greenhouse at an atmospheric $\text{CO}_2$ concentration ( $C_a$ ) of $\approx 450$ ppm and then moved to growth chambers. After a day in darkness to drain the starch reserves, the plants were grown at different levels of $C_a$ (180, 280, 450, 700, 1500 ppm) corresponding to different levels of intercellular $\text{CO}_2$ concentration ( $C_i = 140, 206, 328, 531$ , and $1365$ ppm) for two days. During gas exchange measurements slightly lower $C_i$ levels prevailed (136, 202, 304, 516, 1282 ppm). Symbols: $A_j$ , modelled RuBP-regeneration-limited $\text{CO}_2$ assimilation; $A_m$ , measured $\text{CO}_2$ assimilation; $R_{a,p}$ , respiration by the plastidial anaplerotic pathway; $R_x$ , day respiration by processes other than the plastidial anaplerotic pathway.
<sup>(a)</sup> $A_j$ estimates based on equation 28 with $R_x = 0.96 \mu\text{mol CO}_2 \text{m}^{-2} \text{s}^{-1}$ .
<sup>(b)</sup> Estimates based on gas exchange data (Eqn. 13).
<sup>(c)</sup> Probably overestimated (see text).
<sup>(d)</sup> Estimates based on previously published isotope data (Wieloch, 2022).
<sup>(e)</sup> Estimates based on equation 28 for $C_a \approx 180, 280$ , and $450$ ppm, and equation 29 for $C_a \approx 700$ , and $1500$ ppm.

**Table 3.**
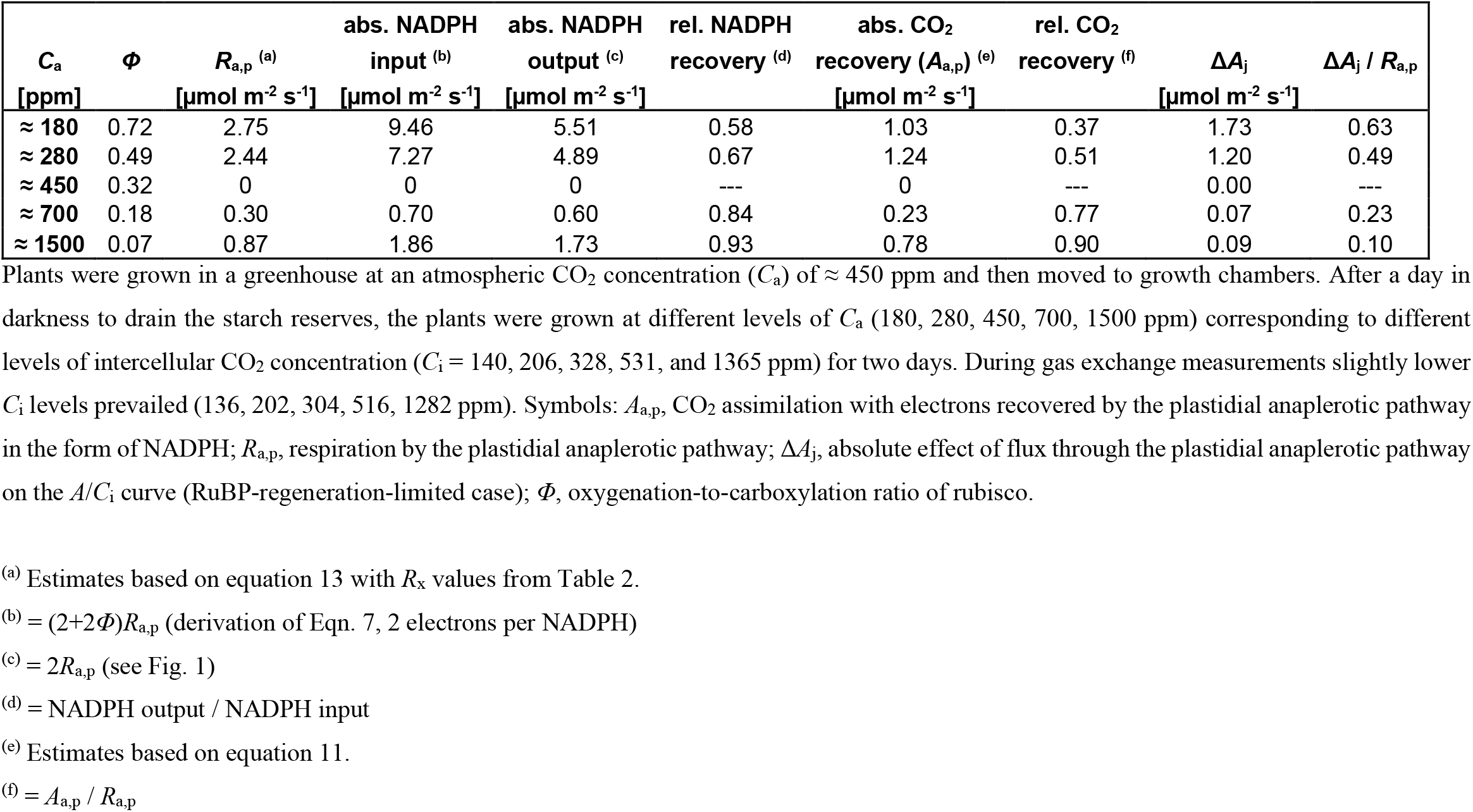
Energy recovery by the plastidial anaplerotic pathway and associated CO_2_ assimilation in sunflower leaves.

| $C_a$ | $\Phi$ | $R_{a,p}$ <sup>(a)</sup> | abs. NADPH<br>input <sup>(b)</sup> | abs. NADPH<br>output <sup>(c)</sup> | rel. NADPH<br>recovery <sup>(d)</sup> | abs. CO<sub>2</sub><br>recovery ($A_{a,p}$) <sup>(e)</sup> | rel. CO<sub>2</sub><br>recovery <sup>(f)</sup> | $\Delta A_j$ | $\Delta A_j / R_{a,p}$ |
| --- | --- | --- | --- | --- | --- | --- | --- | --- | --- |
| [ppm] | | [$\mu\text{mol m}^{-2} \text{s}^{-1}$] | [$\mu\text{mol m}^{-2} \text{s}^{-1}$] | [$\mu\text{mol m}^{-2} \text{s}^{-1}$] | | [$\mu\text{mol m}^{-2} \text{s}^{-1}$] | | [$\mu\text{mol m}^{-2} \text{s}^{-1}$] | |
| $\approx 180$ | 0.72 | 2.75 | 9.46 | 5.51 | 0.58 | 1.03 | 0.37 | 1.73 | 0.63 |
| $\approx 280$ | 0.49 | 2.44 | 7.27 | 4.89 | 0.67 | 1.24 | 0.51 | 1.20 | 0.49 |
| $\approx 450$ | 0.32 | 0 | 0 | 0 | --- | 0 | --- | 0.00 | --- |
| $\approx 700$ | 0.18 | 0.30 | 0.70 | 0.60 | 0.84 | 0.23 | 0.77 | 0.07 | 0.23 |
| $\approx 1500$ | 0.07 | 0.87 | 1.86 | 1.73 | 0.93 | 0.78 | 0.90 | 0.09 | 0.10 |

Plants were grown in a greenhouse at an atmospheric CO<sub>2</sub> concentration ( $C_a$ ) of $\approx 450$ ppm and then moved to growth chambers. After a day in darkness to drain the starch reserves, the plants were grown at different levels of $C_a$ (180, 280, 450, 700, 1500 ppm) corresponding to different levels of intercellular CO<sub>2</sub> concentration ( $C_i$ = 140, 206, 328, 531, and 1365 ppm) for two days. During gas exchange measurements slightly lower $C_i$ levels prevailed (136, 202, 304, 516, 1282 ppm). Symbols: $A_{a,p}$ , CO<sub>2</sub> assimilation with electrons recovered by the plastidial anaplerotic pathway in the form of NADPH; $R_{a,p}$ , respiration by the plastidial anaplerotic pathway; $\Delta A_j$ , absolute effect of flux through the plastidial anaplerotic pathway on the $A/C_i$ curve (RuBP-regeneration-limited case); $\Phi$ , oxygenation-to-carboxylation ratio of rubisco.
<sup>(a)</sup> Estimates based on equation 13 with $R_x$ values from Table 2.
<sup>(b)</sup> = $(2+2\Phi)R_{a,p}$ (derivation of Eqn. 7, 2 electrons per NADPH)
<sup>(c)</sup> = $2R_{a,p}$ (see Fig. 1)
<sup>(d)</sup> = NADPH output / NADPH input
<sup>(e)</sup> Estimates based on equation 11.
<sup>(f)</sup> = $A_{a,p} / R_{a,p}$

**Figure 3.**
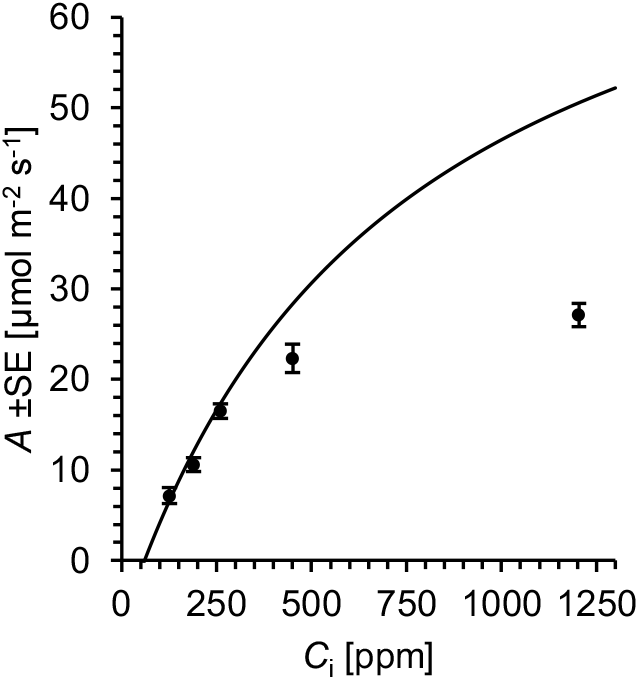
Net CO_2_ assimilation (*A*) of sunflower leaves as function of intercellular CO_2_ concentration (*C*_i_). Black dots: measured values (*A*_m_, *n* = 8). Solid line: Best fit between the data at *C*_a_ = 173, 269, 432 ppm and a model of rubisco-limited *A* (Eqn. 5). Plants were raised in a greenhouse at an atmospheric CO_2_ concentration (*C*_a_) of ≈ 450 ppm and a light intensity of 300-400 μmol photons m^−2^ s^−1^. Gas exchange measurements were performed at 22 °C, 900 μmol photons m^−2^ s^−1^, and atmospheric CO_2_ concentrations of *C*_a_ = 173, 269, 432, 674, 1430 ppm corresponding to intercellular CO_2_ concentrations of *C*_i_ = 126, 188, 264, 450, 1204 ppm. Model inputs: *C*_c_ = 111, 168, 226 μbar, *K*_c_ = 447 μbar CO_2_, *K*_o_ = 439 mbar O_2_, and *Γ*\* = 45.3 μbar CO_2_, *O*_c_ = 206373 μbar, *R*_a,p_ + *R*_x_ = 0.83 μmol CO_2_ m^−2^ s^−1^. Model output: *V*_cmax_ = 83.2 μmol m^−2^ s^−1^.

Based on Farquhar and Wong (1984), *J* is given as

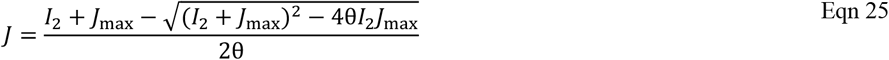

where θ denotes the empirical curvature factor of the light response of electron transport commonly set to 0.7 (Evans, 1989), *J*_max_ denotes the light-saturated rate of linear electron transport, and *I*_2_ denotes incident quanta utilised in electron transport through photosystem II given as

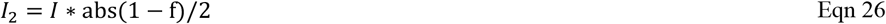

where *I* denotes incident photon flux, abs denotes absorptance of leaves commonly set to 0.85, and f denotes a correction factor for the spectral quality of light commonly set to 0.15 (Evans, 1987). Since gas exchange data collected at very high irradiance are unavailable, *J*_max_ was estimated based on its close relationship with *V*_cmax_ according to published procedures (Walker *et al*., 2014) as

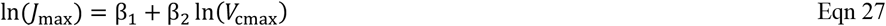

Using parameter estimates pertaining to data reported by Wullschleger (1993) (β_1_ = 1.425, β_2_ = 0.837, *R*^*2*^ = 87.2, p < 0.001, n = 110), we estimate *J*_max_ = 168.3 μmol m^−2^ s^−1^.

## Results

### Metabolic phases of the *A*/*C*_i_ curve

*A*/*C*_i_ curves enable identification of metabolic processes limiting net CO_2_ assimilation (Farquhar *et al*., 1980). Initial linear increases at low *C*_i_ are commonly attributed to rubisco-limited assimilation. A change point in the response of *A* to *C*_i_ marks the onset of RuBP-regeneration-limited assimilation. Furthermore, under high light, *A*/*C*_i_ curves may level out at high *C*_i_ which is commonly attributed to limited triose phosphate utilisation (Sharkey, 1985).

Despite the relatively low resolution of our *A*/*C*_i_ curve, two phases are evident (Fig. 4a). An initial linear increase (dashed line) is followed by RuBP-regeneration-limited assimilation whereas levelling out at high *C*_i_ is not evident. Hence, triose phosphate utilisation did not affect *A*_m_ (denoting measured net CO_2_ assimilation). This is as expected considering the low irradiance applied during data collection (300 μmol photons m^−2^ s^−1^).

**Figure 4.**
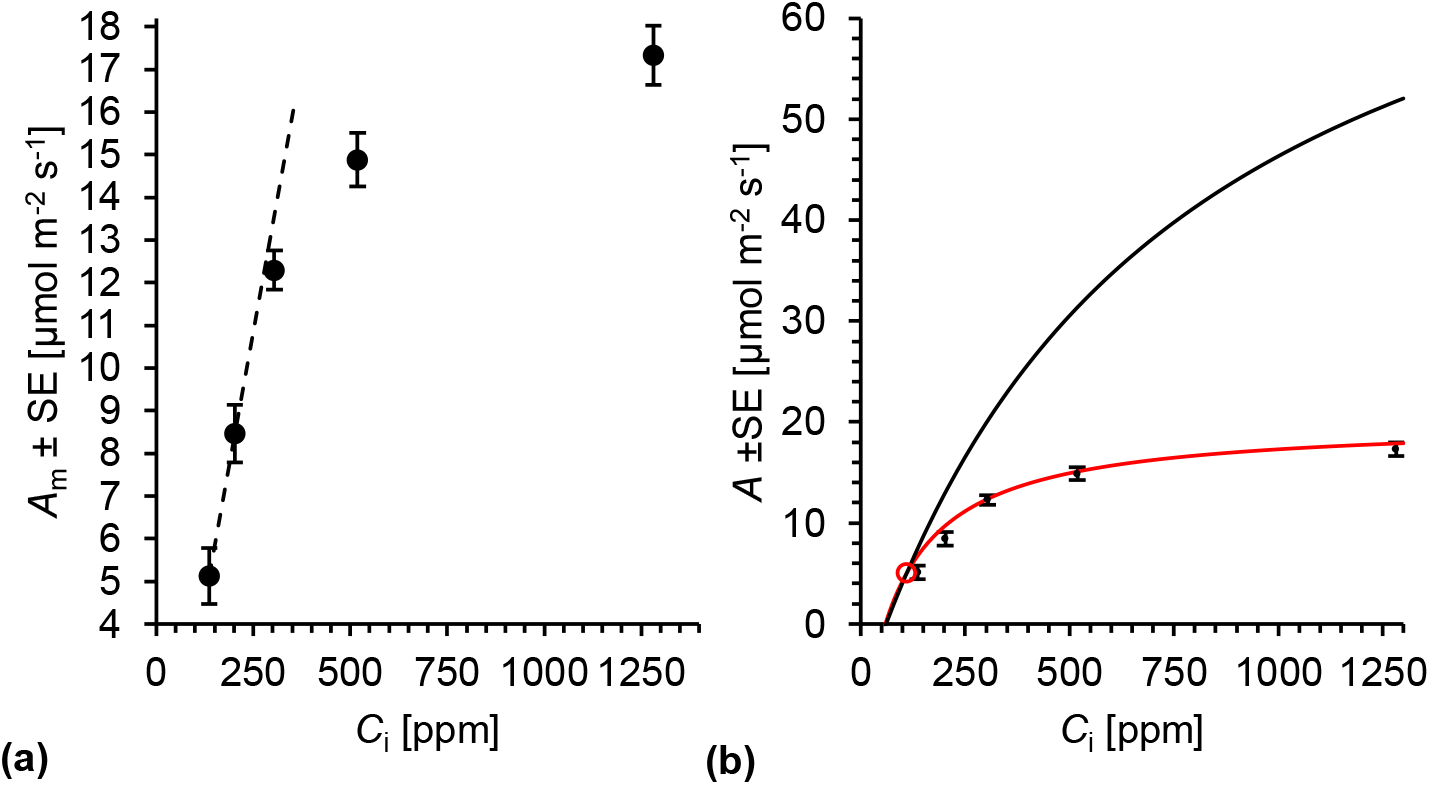
Measured **(a)** and modelled **(b)** net CO_2_ assimilation (*A*) of sunflower leaves as function of intercellular CO_2_ concentration (*C*_i_). Black dots: measured values (*A*_m_, *n* = 8). Dashed black line: Trendline passing through the two lowest *A*_m_ values. Solid black and red lines: modelled rubisco-limited assimilation (*A*_c_) and RuBP-regeneration-limited assimilation (*A*_j_), respectively (respiration by the plastidial anaplerotic pathway not considered). Red circle: intersection between *A*_c_ and *A*_j_. Plants were grown in a greenhouse at an atmospheric CO_2_ concentration (*C*_a_) of ≈ 450 ppm and then moved to growth chambers. After a day in darkness to drain the starch reserves, plants were grown at different levels of *C*_a_ (180, 280, 450, 700, 1500 ppm) corresponding to different levels of *C*_i_ (140, 206, 328, 531, and 1365 ppm) for two days. During gas exchange measurements slightly lower *C*_i_ levels prevailed (136, 202, 304, 516, 1282 ppm). *A*_j_-model inputs: *Γ*\* = 45.3 μbar CO_2_, *J*_max_ = 168.3 μmol m^−2^ s^−1^, θ = 0.7, *I* = 300 μmol photons m^−2^ s^−1^, abs = 0.85, and f = 0.15. *A*_j_-model output: *R*_a,p_ + *R*_x_ = 0.96 μmol CO_2_ m^−2^ s^−1^. *A*_c_-model inputs: *K*_c_ = 447 μbar CO_2_, *K*_o_ = 439 mbar O_2_, and *Γ*\* = 45.3 μbar CO_2_, *O*_c_ = 207548 μbar, *V*_cmax_ = 83.2 μmol m^−2^ s^−1^, *R*_a,p_ + *R*_x_ = 0.96 μmol CO_2_ m^−2^ s^−1^.

Previously, we have shown that flux through the plastidial anaplerotic pathway is not significantly different from zero at *C*_a_ ≈ 450 ppm (Wieloch *et al*., 2022a; Wieloch, 2022). Hence, RuBP-regeneration-limited assimilation for this datapoint as derived from equation 12 is given as

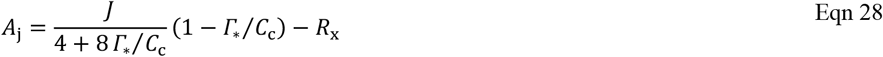

which is equivalent to the original FvCB model. Equation 28 was fitted to *A*_m_ by varying *R*_x_ until the offset between modelled and measured ≈ 450 ppm values was zero (Fig. 4b, red line). The condition was met at *R*_x_ = 0.96 μmol CO_2_ m^−2^ s^−1^. Hence, at ≈ 450 ppm, *R*_d_ proceeds at 7.8% relative to *A*_m_ which agrees well with previous reports. Under standard conditions (400 ppm CO_2_, 21% O_2_, 20-25 °C), leaf *R*_d_ commonly proceeds at ≥ 5% relative to *A*_m_ (Tcherkez *et al*., 2017). This agreement lends some credibility to the validity of our chosen model parameterisation.

Using *R*_x_ = 0.96 μmol CO_2_ m^−2^ s^−1^ and assuming *R*_a,p_ = 0, we next modelled rubisco-limited assimilation based on equation 5 (Fig. 4b, black line). Above *C*_a_ ≈ 133 ppm (*C*_i_ ≈ 110 ppm), we found that *A*_c_ > *A*_j_ ≥ *A*_m_ (Fig. 4b, black line versus red line versus black dots). This indicates that *A*_m_ at *C*_a_ > 133 ppm does not result from rubisco-limited but RuBP-regeneration-limited assimilation.

### Effects of varying rates of day respiration on the *A*/*C*_i_ curve

Based on changes in hydrogen isotope fractionation in starch extracted from the leaves studied here, we previously estimated *R*_a,p_ proceeds at > 7%, > 5%, 0%, ≈ 2%, and ≈ 5% relative to *A*_m_ at *C*_a_ ≈ 180, 280, 450, 700, and 1500 ppm, respectively (Table 2) (Wieloch, 2022). That is, *R*_a,p_ exhibits a change point in response to *C*_a_ at *C*_a_ ≈ 450 ppm. Seeing that there are positive offsets between *A*_m_ and *A*_j_ both above and below *C*_a_ ≈ 450 ppm (Table 2; Fig. 4b, red line versus black dots), this same change point appears to be present in *A*_m_, and *R*_a,p_ can be expected to contribute to the offsets.

Using equation 13, we estimate *R*_a,p_ proceeds at 54%, 29%, 0%, 5%, and 29% relative to *A*_m_ at *C*_a_ ≈ 180, 280, 450, 700, and 1500 ppm, respectively (Table 2). Estimates at low *C*_a_ (≈ 180, and 280 ppm) are consistent with isotope-derived estimates because, based on properties of the biochemical system, the isotope approach can be expected to underestimates *R*_a,p_ at low *C*_a_ (see Wieloch *et al*., 2022a). By contrast, the isotope approach is believed to not strongly underestimate *R*_a,p_ at high *C*_a_ (Wieloch, 2022). However, gas-exchange estimates of *R*_a,p_ at high *C*_a_ (≈ 700, and 1500 ppm) are considerably larger than isotope-derived estimates. This may suggest that gas exchange modelling overestimates *R*_a,p_ at high *C*_a_.

Overestimation of *R*_a,p_ at high *C*_a_ may derive from variability of *R*_x_. To follow up on this, we solve the *A*_j_ model (Eqn. 12) for *R*_x_ and substitute *A*_j_ by *A*_m_ as

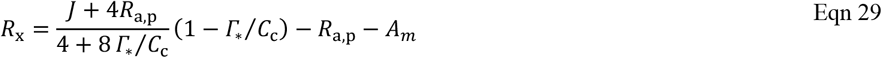

Using isotope-derived *R*_a,p_ values as inputs, we estimate that absolute rates of *R*_x_ increase from 450 to 1500 ppm while relative rates remain approximately constant (Table 2). Based on regulatory properties of leaf respiratory processes, this finding seems reasonable (see ‘Metabolic origin of day respiration at high *C*_a_’). Taken together, our results suggest *A*_m_ is a function of RuBP-regeneration-limited assimilation and variability of *R*_a,p_ and *R*_x_ (Eqn. 12). Figure 5 summarises estimated carbon and associated NADPH fluxes. Uncertainties associated with these estimates are discussed below (‘Model criticism and future research directions’).

**Figure 5.**
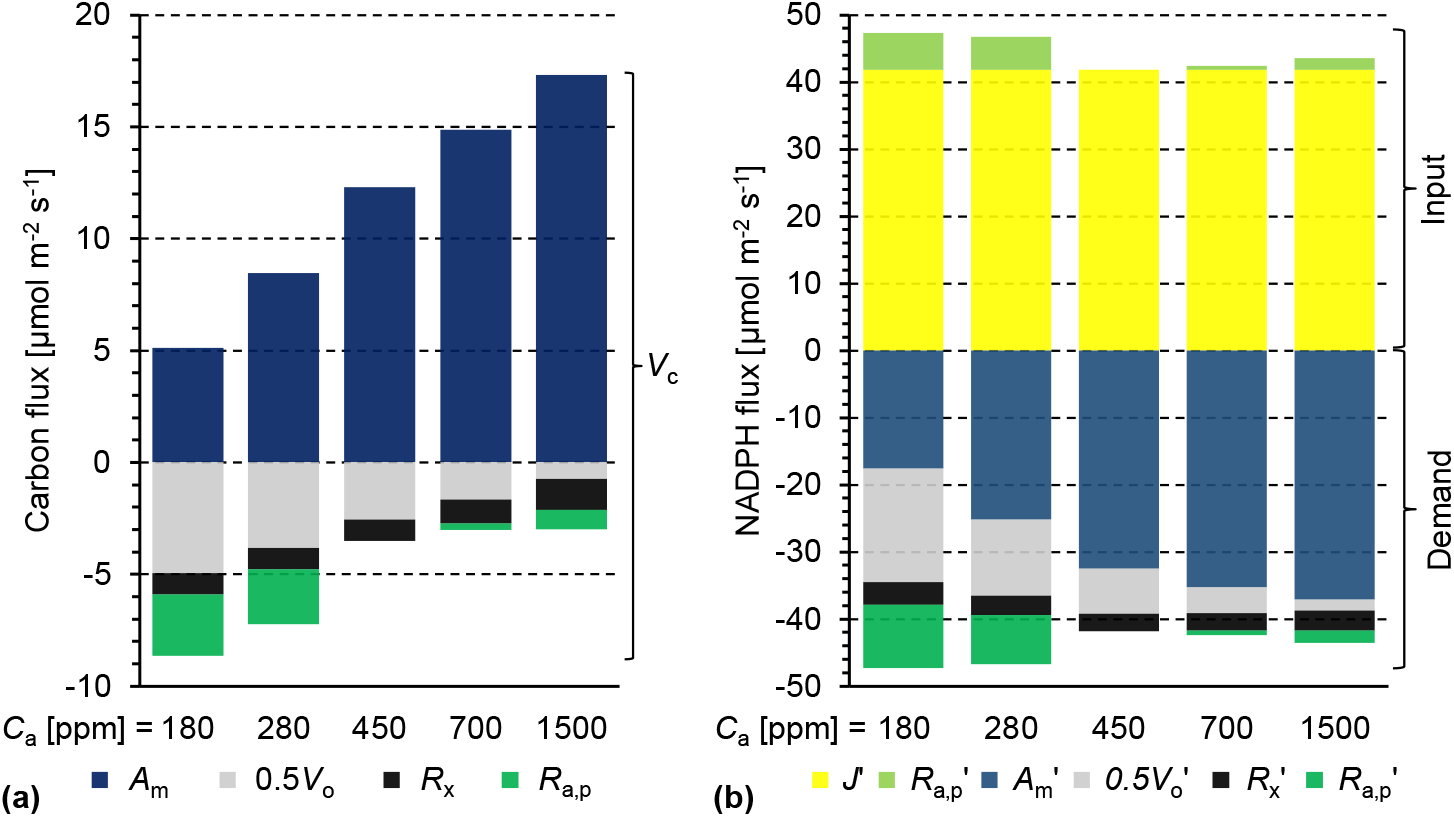
Carbon **(a)** and NADPH fluxes **(b)** at varying atmospheric CO_2_ concentration (*C*_a_) in sunflower leaves. Estimates derived from modelling RuBP-regeneration-limited CO_2_ assimilation. The model considers the Calvin-Benson cycle, the photorespiration cycle, the plastidial anaplerotic pathway, and respiration by processes other than the plastidial anaplerotic pathway. Plants were grown in a greenhouse at *C*_a_ ≈ 450 ppm and then moved to growth chambers. After a day in darkness to drain the starch reserves, plants were grown at different levels of *C*_a_ (180, 280, 450, 700, 1500 ppm) corresponding to different levels of *C*_i_ (140, 206, 328, 531, and 1365 ppm) for two days. During gas exchange measurements slightly lower *C*_i_ levels prevailed (136, 202, 304, 516, 1282 ppm). NADPH demands of individual carbon fluxes (see Fig. 5a) were calculated as (2 + 2*Φ*) * flux with flux = {-*A*_m_; 0.5*V*_o_; *R*_x_; *R*_a,p_}. Symbols: *A*_m_, measured net CO_2_ assimilation; *A*_m_’, NADPH demand of net CO_2_ assimilation; *J*’, NADPH supply by whole chain electron transport (0.5*J*); *R*_a,p_, respiration by the plastidial anaplerotic pathway; *R*_a,p_’, NADPH supply (light green) and demand (dark green) by the plastidial anaplerotic pathway; *R*_x_, day respiration by processes other than the plastidial anaplerotic pathway; *R*_x_’, NADPH demand for assimilation of CO_2_ respired by processes other than the plastidial anaplerotic pathway; *V*_c_, rubisco carboxylation; 0.5*V*_o_, photorespiratory CO_2_ release; 0.5*V*_o_’, NADPH demand for assimilation of CO_2_ respired by photorespiration; *Φ*, oxygenation-to-carboxylation ratio of rubisco.

### Integration of day respiration in carbon metabolism

Based on estimations and assumptions given above, *R*_a,p_ proceeds at ≈ 54%, 29%, 0%, 2%, and 5% relative to *A*_m_ at *C*_a_ ≈ 180, 280, 450, 700, and 1500 ppm, respectively (Fig. 6a). Thus, it governs *R*_d_ at low *C*_a_ (≈ 73% contribution at ≈ 180, and 280 ppm) and contributes substantially to *R*_d_ at high *C*_a_ (≈ 22% at ≈ 700 ppm, and ≈ 39% at ≈ 1500 ppm; Fig. 6b). Taken together, *R*_d_ proceeds at ≈ 72%, 40%, 8%, 9%, and 13% relative to *A*_m_ at *C*_a_ ≈ 180, 280, 450, 700, and 1500 ppm, respectively (Table 2). While *R*_a,p_ exceeds photorespiratory CO_2_ release at high *C*_a_ (≈ 1500 ppm), it proceeds at > 55% relative to the rate of photorespiratory CO_2_ release at low *C*_a_ (≈ 180, and 280 ppm; Fig. 6c). Furthermore, *R*_a,p_ accounts for > 40% of the total CO_2_ release at both low and high *C*_a_ (≈ 180, 280, and 1500 ppm; Fig. 6d). At low *C*_a_ (≈ 180, and 280 ppm), *R*_a,p_ accounts for > 15% of *V*_c_ (Fig. 6e) and thus causes significant futile carbon cycling involving CO_2_ uptake by the CBC and release by the plastidial anaplerotic pathway. At the same *C*_a_, > 10% of all RuBP is provided by the plastidial anaplerotic pathway (Fig. 6f). When plotted against *A*_m_, *R*_a,p_ increases nonlinearly both below and above *C*_a_ ≈ 450 ppm (Fig. 6g). Towards very low *A*_m_ (< 5 μmol m^−2^ s^−1^, *C*_a_ < 180 ppm), *R*_a,p_ may approach a maximum. Above *A*_m_ = 12.3 μmol m^−2^ s^−1^ (*C*_a_ ≈ 450 ppm), *R*_a,p_ increases exceed *R*_x_ increases by approximately a factor of two (Fig. 6g-h).

**Figure 6.**
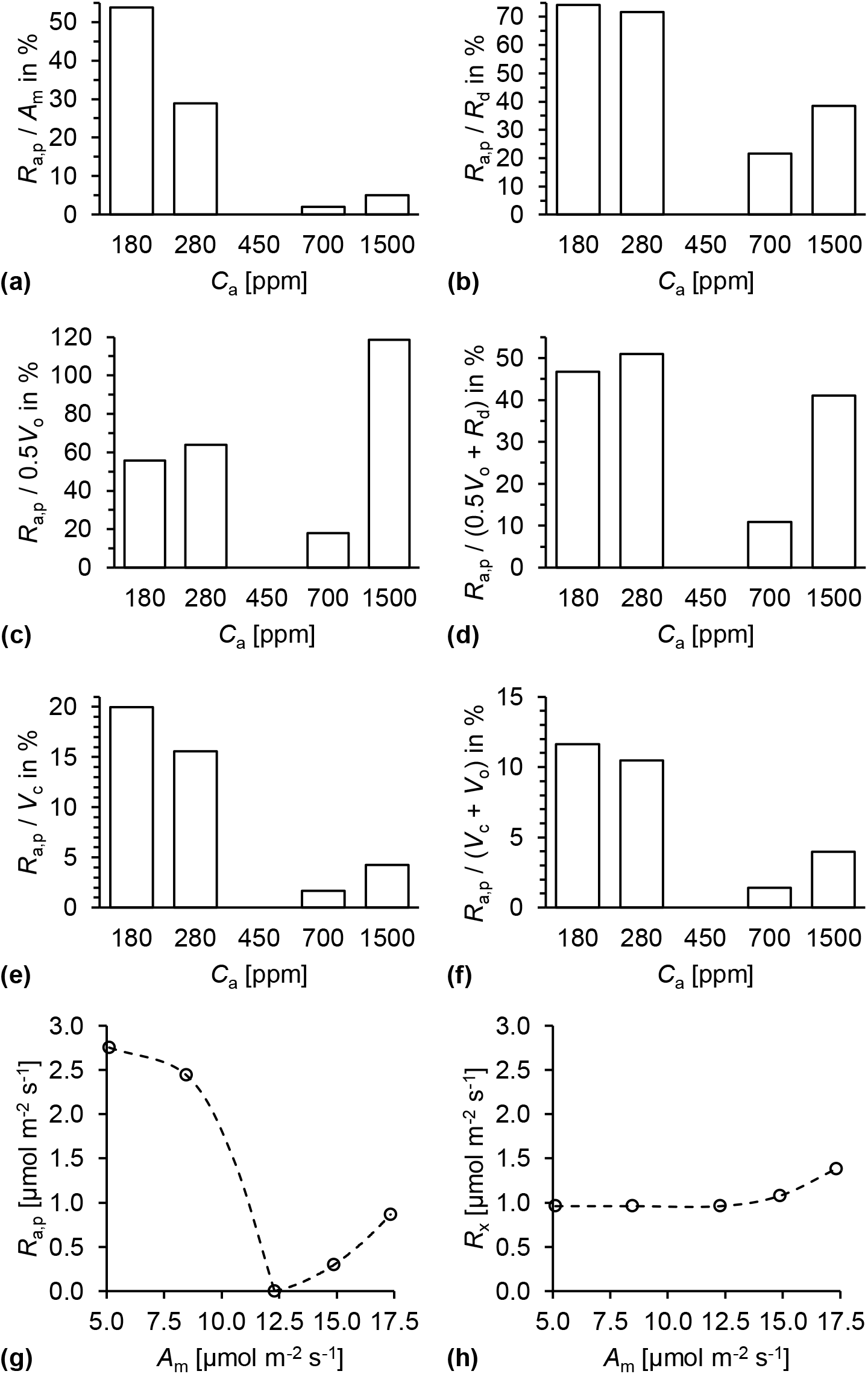
Integration of day respiration in carbon metabolism of sunflower leaves. Plants were grown in a greenhouse at *C*_a_ ≈ 450 ppm and then moved to growth chambers. After a day in darkness to drain the starch reserves, plants were grown at different levels of *C*_a_ (180, 280, 450, 700, 1500 ppm) corresponding to different levels of *C*_i_ (140, 206, 328, 531, and 1365 ppm) for two days. During gas exchange measurements slightly lower *C*_i_ levels prevailed (136, 202, 304, 516, 1282 ppm). Symbols: *A*_m_, measured CO_2_ assimilation; *C*_a_, atmospheric CO_2_ concentration; *C*_i_, intercellular CO_2_ concentration; *R*_a,p_, respiration by and flux through the plastidial anaplerotic pathway; *R*_d_, total day respiration; *R*_x_, day respiration by processes other than the plastidial anaplerotic pathway; *V*_c_, rubisco carboxylation; *V*_o_, rubisco oxygenation. All panels show discrete data. Dashed lines in added to guide the eye.

### Energy recovery by the plastidial anaplerotic pathway and associated CO_2_ assimilation

NADPH inputs required to assimilate CO_2_ respired by the plastidial anaplerotic pathway exceed NADPH outputs of this pathway (Table 3). However, relative NADPH loss decreases as *Φ* decreases with relative NADPH recovery converging to one. Accordingly, CO_2_ release by the anaplerotic pathway (*R*_a,p_) exceeds CO_2_ recovery with NADPH from the anaplerotic pathway (*A*_a,p_, Table 3). Relative CO_2_ recovery increases with decreasing *Φ* converging to one (*A*_a,p_ / *R*_a,p_). Hence, *R*_a,p_ is larger than the effect that flux through the plastidial anaplerotic pathway has on *A*/*C*_i_ curves (Δ*A*_j_, Table 3). With decreasing *Φ*, the relative effect of flux through the plastidial anaplerotic pathway on the *A*/*C*_i_ curve decreases converging to zero (Δ*A*_j_ / *R*_a,p_, Table 3, black circles in Fig. 2). Taken together, anaplerotic flux is energetically wasteful and thus decreases net CO_2_ assimilation but this is partly offset by NADPH recovery. Recovered NADPH is most efficiently used for CO_2_ recovery when photorespiration is low since photorespiration is energetically wasteful and releases additional CO_2_ (see ‘Theory’).

### Effects of the plastidial anaplerotic pathway on the light reactions of photosynthesis

In the present study, light was maintained at 300 μmol photons m^−2^ s^−1^ across all *C*_a_ treatments. Accordingly, linear electron transport supplied a steady amount of NADPH (Fig. 5b, yellow bars). Above and below *C*_a_ ≈ 450 ppm, the plastidial anaplerotic pathway supplied additional NADPH yet no ATP (Fig. 5b, light green bars). Balancing the ensuing NADPH surplus with the demands of the CBC and photorespiration requires additional inputs of ATP from photophosphorylation (Fig. 7, black triangles). Hence, the ratio of ATP-to-NADPH demand from the light reactions increases both above and below *C*_a_ ≈ 450 ppm with increasing flux through the plastidial anaplerotic pathway (Fig. 7, red circles).

**Figure 7.**
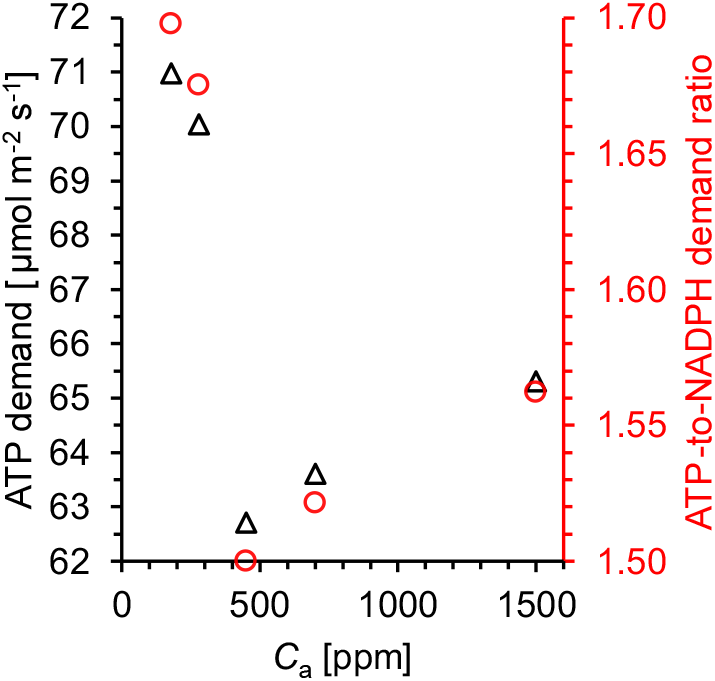
Energy demand from the light reactions of photosynthesis at varying atmospheric CO_2_ concentration (*C*_a_) in sunflower leaves. Black triangles: ATP demand. Red circles: ATP-to-NADPH demand ratio. Energy fluxes were estimated based on carbon flux estimates from modelling RuBP-regeneration-limited CO_2_ assimilation. The model considers the Calvin-Benson cycle, the photorespiration cycle, the plastidial anaplerotic pathway, and respiration by processes other than the plastidial anaplerotic pathway. Plants were grown in a greenhouse at *C*_a_ ≈ 450 ppm and then moved to growth chambers. After a day in darkness to drain the starch reserves, plants were grown at different levels of *C*_a_ (180, 280, 450, 700, 1500 ppm) corresponding to different levels of *C*_i_ (140, 206, 328, 531, and 1365 ppm) for two days. During gas exchange measurements slightly lower *C*_i_ levels prevailed (136, 202, 304, 516, 1282 ppm).

Assuming the synthesis of 3 mol ATP requires 12 mol H^+^ as derived from the thermodynamics of the ATPase reaction (Steigmiller *et al*., 2008), we find that ATP-to-NADPH supply by linear electron transport including Q cycling is perfectly aligned with the demands of the CBC and photorespiration at *C*_a_ ≈ 450 ppm (Table 4, *cf*. Eqn. 22). By contrast, assuming the synthesis of 3 mol ATP requires 14 mol H^+^ as derived from ATPase subunit composition (Seelert *et al*., 2000), linear electron transport including Q cycling results in an ATP deficit at *C*_a_ ≈ 450 ppm. In both cases, ATP deficits increase significantly both above and below *C*_a_ ≈ 450 ppm. This suggests that increasing flux through the plastidial anaplerotic pathway increases the requirement for ATP synthesis via cyclic electron transport around photosystem I. Assuming an H^+^-ATP synthesis ratio of 12/3, cyclic electron transport including Q cycling is not required when the plastidial anaplerotic pathway is inactive, but it may account for up to 17% of the rate of whole electron transport when this pathway is active. Assuming an H^+^-ATP synthesis ratio of 14/3, cyclic electron transport including Q cycling accounts for ≈ 20% of the rate of whole electron transport when the plastidial anaplerotic pathway is inactive and may increase to above 30% when this pathway is active. Thus, flux through the plastidial anaplerotic pathway may affect the light reactions of photosynthesis.

**Table 4.** Effects of the plastidial anaplerotic pathway on electron transport in sunflower leaves.

| H<sup>+</sup>-ATP ratio of ATPase | 12/3 |  |  |  |  | 14/3 |  |  |  |  |
| --- | --- | --- | --- | --- | --- | --- | --- | --- | --- | --- |
| C<sub>a</sub> [ppm] | 180 | 280 | 450 | 700 | 1500 | 180 | 280 | 450 | 700 | 1500 |
| LET [$\mu\text{mol m}^{-2} \text{s}^{-1}$] | 83.6 | 83.6 | 83.6 | 83.6 | 83.6 | 83.6 | 83.6 | 83.6 | 83.6 | 83.6 |
| ATP demand [$\mu\text{mol m}^{-2} \text{s}^{-1}$] | 71.0 | 70.0 | 62.7 | 63.6 | 65.3 | 71.0 | 70.0 | 62.7 | 63.6 | 65.3 |
| ATP from LET [$\mu\text{mol m}^{-2} \text{s}^{-1}$] | 62.7 | 62.7 | 62.7 | 62.7 | 62.7 | 53.8 | 53.8 | 53.8 | 53.8 | 53.8 |
| ATP deficit [$\mu\text{mol m}^{-2} \text{s}^{-1}$] | 8.3 | 7.3 | 0.0 | 0.9 | 2.6 | 17.2 | 16.3 | 9.0 | 9.9 | 11.6 |
| CET [$\mu\text{mol m}^{-2} \text{s}^{-1}$] | 16.5 | 14.7 | 0.0 | 1.8 | 5.2 | 40.2 | 38.0 | 20.9 | 23.0 | 27.0 |
| CET/(CET+LET) | 0.17 | 0.15 | 0.0 | 0.02 | 0.06 | 0.32 | 0.31 | 0.20 | 0.22 | 0.24 |

Plants were grown in a greenhouse at an atmospheric CO<sub>2</sub> concentration (C<sub>a</sub>) of ≈ 450 ppm and then moved to growth chambers. After a day in darkness to drain the starch reserves, the plants were grown at different levels of C<sub>a</sub> (180, 280, 450, 700, 1500 ppm) for two days. Gas exchange measurements were performed at a light intensity of 300 $\mu\text{mol photons m}^{-2} \text{s}^{-1}$ . Estimations pertaining to linear and cyclic electron transport (LET and CET, both including Q cycling) assumed transmembrane transport of 3 and 2 H<sup>+</sup> per electron, respectively. Synthesis of 3 mol ATP by ATPase was assumed to require either 14 or 12 mol H<sup>+</sup> (Seelert *et al.*, 2000; Steigmiller *et al.*, 2008). Processes not considered include the water-water cycle (affects the ATP-to-NADPH supply ratio of electron transport), GAP/3PGA cycles and the chloroplast malate valve (affect ATP-to-NADPH demand ratios of carbon metabolism).

## Discussion

Here, we expanded FvCB models by terms accounting for respiration and energy recycling by the plastidial anaplerotic pathway (Eqs. 5, and 12). We fitted the model of RuBP-regeneration-limited assimilation to gas exchange data of sunflower leaves by adjusting day respiration. This approach and previous reports (Wieloch *et al*., 2022a; Wieloch, 2022) challenge a longstanding FvCB modelling assumption, namely, the treatment of day respiration as independent, constant term. If this assumption is false, then several FvCB-model-based methods require revision, including methods for estimating day respiration (e.g., Laisk, 1977) and mesophyll conductance (see below). Moreover, initial linear increases in *A*/*C*_i_ curves are commonly interpreted in terms of rubisco-limited assimilation (Farquhar *et al*., 1980). Here, we propose RuBP-regeneration-limited assimilation and varying rates of respiration from the plastidial anaplerotic pathway as alternative explanation. Our modelling results suggest that the plastidial anaplerotic pathway can be an important player in plant carbon and energy metabolism affecting properties such as the rate of day respiration, net CO_2_ assimilation, net carboxylation, net oxygenation (photorespiration), RuBP regeneration, NADPH availability, the ATP-to-NADPH demand ratio of plastidial carbon metabolism, and probably the contribution of the cyclic electron pathway to supplying ATP (Figs. 5-7, Tables 2-4).

### Consistent evidence in support of flux through the plastidial anaplerotic pathway

According to our gas exchange analysis, flux through the plastidial anaplerotic pathway proceeds at 0, 2.44, and 2.75 μmol m^−2^ s^−1^ at *C*_a_ ≈ 450, 280, and 180 ppm, respectively (Table 2, Fig. 5a). That is, flux increases with decreasing *C*_a_ from ≈ 0% over ≈ 29% to ≈ 54% relative to the rate of net CO_2_ assimilation (Fig. 6a). This is consistent with regulatory properties of the pathway and previously reported flux estimates as discussed next.

Flux through the plastidial anaplerotic pathway is controlled at its first enzyme, glucose-6-phosphate dehydrogenase (G6PD, Fig. 1). Under illumination, G6PD is downregulated by a thioredoxin dependent mechanism (Née et al., 2009). However, downregulation can be reversed allosterically by G6P (Cossar *et al*., 1984; Preiser *et al*., 2019). Under medium to high *C*_a_, the reaction catalysed by plastidial phosphoglucose isomerase (PGI) was found to be removed from equilibrium on the side of F6P (Schleucher *et al*., 1999; Wieloch *et al*., 2022a; Wieloch, 2022) resulting in low G6P concentration (Dietz, 1985; Gerhardt *et al*., 1987; Kruckeberg *et al*., 1989). With shifts to low *C*_a_, the PGI reaction moves towards equilibrium (Wieloch *et al*., 2022a; Wieloch, 2022) and plastidial G6P concentrations increase (Dietz, 1985). Furthermore, phosphorolytic starch breakdown increases with photorespiration providing additional G6P (dotted arrow in Fig. 1; Weise *et al*., 2006). These mechanisms are consistent with reports of negligible flux through the anaplerotic pathway at medium *C*_a_ (low G6P concentration) and flux increases with decreasing *C*_a_, increasing photorespiration, and increasing drought (high G6P concentration; Wieloch *et al*., 2018, 2022a,b; Wieloch, 2022).

Previously, we estimated flux through the plastidial anaplerotic pathway in the leaves studied here proceeds at 0%, > 5% and > 7% relative to the rate of net CO_2_ assimilation at *C*_a_ ≈ 450, 280, and 180 ppm, respectively (Wieloch *et al*., 2022a; Wieloch, 2022). These estimates are based on a hydrogen isotope signal in starch introduced in the starch biosynthesis pathway at the level of G6P H^1^. G6P may be converted back to fructose 6-phosphate (F6P) by PGI, and F6P may leave the starch biosynthesis pathway via transketolase. Combinedly, G6P to F6P conversion and F6P use by transketolase can be expected to cause washout of the isotope signal from the starch biosynthesis pathway. As evident from an isotope signal at starch glucose H^2^, the plastidial PGI reaction is strongly removed from equilibrium on the side of F6P at *C*_a_ ≥ 450 ppm (Wieloch *et al*., 2022a; Wieloch, 2022). This is thought to impede back conversion of G6P to F6P and concomitant signal washout. However, the PGI reaction moves progressively towards equilibrium as *C*_a_ decreases below ≈ 450 ppm (Wieloch *et al*., 2022a; Wieloch, 2022). This is thought to result in increasing back conversion of G6P to F6P and signal washout. Thus, increasing offsets between *R*_a.p_ estimates from isotope and gas exchange analysis with decreasing *C*_a_ probably derive from increasing washout of the isotope signal.

### Physiological function of flux through the plastidial anaplerotic pathway

Flux through the anaplerotic pathway lowers net CO_2_ assimilation below theoretically possible values (Fig. 4b, black dots versus red line). As a result of this flux, sunflower leaves merely achieve ≈ 88% and ≈ 75% of their theoretical assimilatory potential at *C*_a_ ≈ 280 ppm and ≈ 180 ppm, respectively. Similarly, the anaplerotic pathway recovers only part of the energy it consumes (Fig. 5b, Table 3), and energy dissipation is surely not physiologically beneficial under RuBP-regeneration-limited conditions. Thus, for the conditions and period studied here, the anaplerotic pathway appears to be detrimental for both leaf carbon and energy balances.

However, the sunflowers studied here were raised at *C*_a_ ≈ 450 ppm. Hence, heterotrophic carbon demands were probably adapted to carbon inputs at *C*_a_ ≈ 450 ppm. Moving the plants into low-*C*_a_ environments may have caused divergences from the previously established whole-plant steady state with leaf carbon exports exceeding net CO_2_ assimilation (Fig. 1). This source limitation may have resulted in excess export of triose phosphates from chloroplasts. Plastidial triose phosphate shortage would impede RuBP regeneration. To maintain the supply of RuBP, plants may inject G6P from phosphorolytic starch breakdown into the CBC (Weise *et al*., 2006). G6P-derived carbon can enter the CBC via transketolase without respiratory carbon loss and via the anaplerotic pathway with respiratory carbon loss. However, transketolase and other enzymes in the regeneration part of the CBC require triose phosphate. When RuBP regeneration is impeded by triose phosphate shortage, the anaplerotic pathway may help to ensure RuBP supply.

RuBP may undergo either oxygenation or carboxylation at rubisco. With decreasing *C*_a_, oxygenation becomes substantial (Table 3, ≈ 1 and ≈ 1.5 mol RuBP oxygenation per 2 mol RuBP carboxylation at *C*_a_ ≈ 280 ppm and ≈ 180 ppm, respectively). Thus, much of the anaplerotically supplied RuBP feeds into photorespiration. As proposed previously, maintaining photorespiration through inputs of RuBP might make sense physiologically in the context of the concept of photorespiration-linked *de novo* nitrogen assimilation (Wieloch *et al*., 2022a). Reportedly, photorespiration rate correlates positively with the rate of *de novo* nitrate assimilation (Bloom, 2015), and it seems plausible that plants would shift metabolism towards nitrogen assimilation when carbon assimilation is impeded (at low *C*_a_). Assimilated nitrogen may support *de novo* protein synthesis which may result in increased photosynthetic capacity at the enzyme level and help to overcome source limitations.

In this context, it is interesting to note that mutants incapable of phosphorolytic starch breakdown show severe phenotypes especially under drought (dotted arrow in Fig. 1; Zeeman *et al*., 2004). This was explained by the loss of G6P injection from starch into the CBC at night (Zeeman *et al*., 2004). Based on findings presented here and previously, impaired G6P injection during the day may contribute to this phenotype (Wieloch *et al*., 2018, 2022a,b). Lastly, G6P injection from starch into the CBC by the anaplerotic pathway may support photosynthetic induction (see below).

### Metabolic origin of day respiration at high *C*_a_

In our gas exchange analysis, respiration by the plastidial anaplerotic pathway was fixed at 0%, 2%, and 5% relative to the rate of net CO_2_ assimilation at *C*_a_ ≈ 450, 700, and 1500 ppm, respectively (Fig. 5a, Table 2). These *R*_a,p_ estimates come from a previously published isotope analysis of the samples studied here (Wieloch, 2022). Increases in *R*_a,p_ above *C*_a_ ≈ 450 ppm are consistent with regulatory properties of the plastidial anaplerotic pathway (Wieloch, 2022). Towards high *C*_a_, plastidial G6P concentrations increase with net CO_2_ assimilation (Dietz, 1985). This can be expected to cause increased G6PD activity and flux through the plastidial anaplerotic pathway (see above). Note, in contrast to the low-*C*_a_ case (see above), *R*_a,p_ increases at high *C*_a_ are not caused by PGI regulation (Wieloch, 2022).

Here, we estimated that respiration by processes other than the plastidial anaplerotic pathway proceeds at 0.96, 1.08, and 1.38 μmol m^−2^ s^−1^ at *C*_a_ ≈ 450, 700, and 1500 ppm, respectively (Fig. 5a, Table 2). These increases are consistent with regulatory properties of the cytosolic OPPP (Wieloch, 2022) which has recently been identified as a major source of day respiration (Xu *et al*., 2022). Flux through the cytosolic OPPP and associated respiration is controlled at the level of the first OPPP enzyme, G6PD (Fig. 1). Under illumination, cytosolic G6PD activity increases with glucose concentration through *de novo* enzyme synthesis (Hauschild & von Schaewen, 2003). In line with this, respiration at 5 °C was shown to increase with glucose concentration and overall leaf soluble sugar concentration (Tjoelker *et al*., 2009; Wieloch, 2022). Generally, increasing *C*_a_ results in increasing leaf soluble sugar concentration (Ainsworth & Long, 2005) and may thus cause increasing respiration by the cytosolic OPPP.

### Does *g*_m_ vary with *C*_a_?

Using the online carbon isotope discrimination method and/or the variable *J* method (Evans *et al*., 1986; Harley *et al*., 1992), several authors reported nonlinear relationships between *g*_m_ and *C*_a_ with similar patterns across different species including sunflower (Flexas *et al*., 2007; Hassiotou *et al*., 2009; Vrábl *et al*., 2009; Xiong *et al*., 2015). As *C*_a_ increases, *g*_m_ initially increases up to a change point beyond which it decreases again. This response is strikingly similar to the *C*_a_-response of day respiration reported here. As *C*_a_ increases, *R*_a,p_ initially decreases up to a change point beyond which it increases again along with *R*_x_ (Fig. 5a, Table 2). Day respiration and *g*_m_ have opposite effects on net CO_2_ assimilation. That is, both processes could explain offsets between measured and theoretically possible values of net CO_2_ assimilation (Fig. 4b, black dots versus red line). However, changes in day respiration estimated here based on gas exchange analysis are corroborated by results from an independent isotope analysis (Wieloch *et al*., 2022a; Wieloch, 2022). Furthermore, underlying metabolic mechanisms seem straight forward (see above). By contrast, mechanisms underlying the *C*_a_-response of *g*_m_ have remained unclear despite considerable research interest over the past three decades (Nadal *et al*., 2021). Several authors suggested that much of the observed variability of *g*_m_ may be due to methodological shortcomings (Tholen *et al*., 2012; Gu & Sun, 2014; Yin *et al*., 2020; Nadal *et al*., 2021). Our results corroborate this viewpoint. Therefore, we recommend modifying the variable *J* method to allow for variability in day respiration. Furthermore, we recommend modifying the online carbon isotope discrimination method to allow for variability in isotope fractionation of photorespiration and day respiration. Adequate (yet probably challenging) parameterisation of these models may show whether *g*_m_ varies with *C*_a_. Similarly, considering day respiratory variability may advance the discussion about relationships between *g*_m_ and other environmental variables (drought, irradiance, ozone, temperature; see Nadal *et al*., 2021).

### Respiration by the plastidial anaplerotic pathway during photosynthetic induction

Flux through the plastidial anaplerotic pathway and associated respiration depend on G6PD activity. Plastidial G6PD is inhibited by reduced thioredoxin (Née et al., 2009). However, reduction of the plastidial thioredoxin pool after illumination is not instantaneous but builds up over the course of several minutes (Scheibe, 1981). This was shown to result in a gradual activation of plastidial malate dehydrogenase (Scheibe, 1981). Similarly, gradual inhibition of plastidial G6PD can be expected. Thus, flux through the plastidial anaplerotic pathway may persist over the course of several minutes after illumination decreasing from initially higher to lower levels. This would affect day respiration, net CO_2_ assimilation, NADPH supply, the isotopic composition of respired CO_2_, etc. Future studies on the variability of photosynthetic parameters such as *g*_m_ during photosynthetic induction (*cf*. Kaiser *et al*., 2017; Sakoda *et al*., 2021; Liu *et al*., 2022) are encouraged to consider this as a possibility.

### Does day respiration explain part of the *A*/*C*_i_ curve drop at high *C*_a_?

If *R*_d_ is held constant, rubisco-limited and RuBP-regeneration-limited net CO_2_ assimilation modelled by the canonical FvCB equations increase continuously with increasing *C*_a_ (Farquhar *et al*., 1980). However, especially under high light, measured *A*/*C*_i_ curves are frequently seen to level out or even decline at high *C*_a_ (e.g., Sharkey, 1985; Harley & Sharkey, 1991). According to our analysis, day respiration increases from ≈ 8% relative to the rate of net CO_2_ assimilation at *C*_a_ ≈ 450 ppm to ≈ 13% at *C*_a_ ≈ 1500 ppm (Fig. 5a, Table 2). This is consistent with positive responses of *R*_d_ to *C*_a_ reported by some authors (e.g., Wang *et al*., 2001). By contrast, other authors reported *R*_d_ is independent of *C*_a_ (e.g., Ayub *et al*., 2011).

Increases in *R*_d_ are consistent with regulatory properties of both the plastidial and cytosolic OPPP (see above). As *C*_a_ increases from medium to high values, plastidial G6P concentrations and leaf soluble sugar concentrations increase (Dietz, 1985; Ainsworth & Long, 2005). This can be expected to activate both plastidial and cytosolic G6PD (Cossar *et al*., 1984; Hauschild & von Schaewen, 2003; Preiser *et al*., 2019), increase respiration by the plastidial and cytosolic OPPP and explain part of the *A*/*C*_i_ curve drop at high *C*_a_. Furthermore, it may explain part of the lower than expected stimulation of net CO_2_ assimilation by increasing *C*_a_ (Wieloch, 2022). However, high sink strengths (e.g., in vigorously growing plants) may counteract the build-up of leaf soluble sugar concentrations and respiration by the plastidial and cytosolic OPPP. This may explain why some studies found *R*_d_ increases at high *C*_a_ whereas others did not.

### Model criticism and future research directions

Generally, modelling net CO_2_ assimilation by FvCB-type models relies on numerous assumptions (Tcherkez & Limami, 2019), and the present study is no exception. For instance, our models assume that photorespiration is a closed cycle. However, some of the carbon entering photorespiration may actually be diverted into other pathways such as *de novo* nitrogen assimilation (Busch *et al*., 2018). Similarly, in our *A*_j_ model (Eqn. 12), the CBC and photorespiration are assumed to consume all energy. However, part of the available energy can be expected to be utilised by other plastidial processes such as the reduction of nitrite to ammonia (Buchanan *et al*., 2015) or NAD(P)H shuttling to other cell compartments (Scheibe, 2004). Furthermore, our modelling assumes constant absolute *R*_x_ at *C*_a_ ≤ 450 ppm (Table 2). However, leaf soluble sugar concentration may decrease with decreasing *C*_a_ (Ainsworth & Long, 2005). This may cause a reduction in G6PD activity and respiration by the cytosolic OPPP (Hauschild & von Schaewen, 2003). Lastly, our modelling relies on sunflower parameter estimates from the literature. Such estimates can vary strongly among species (Orr *et al*., 2016), and there is a lack of data on parameter variability in sunflower. Considering these uncertainties, results reported here require validation (Supporting Information, Notes S1). Hence, we recommend follow-up studies that combine gas exchange measurements, isotope measurements, and model parameter measurements from the same plants.

That said, our estimate of *R*_x_ at *C*_a_ ≈ 450 ppm is consistent with previously reported estimates (Tcherkez *et al*., 2017). Furthermore, estimates of flux through the plastidial anaplerotic pathway from gas exchange analyses presented here are qualitatively consistent with previously reported isotope-derived estimates (Wieloch *et al*., 2022a; Wieloch, 2022). Lastly, all estimates of day respiration are consistent with regulatory properties of the plastidial and cytosolic anaplerotic pathway. Hence, we hope that the model extensions and ideas presented here will help to advance photosynthesis modelling. Future research directions of interest include the study of *R*_a,p_ at and below the CO_2_ compensation point, and attempts to model *R*_a,p_ as function of underlying biochemical mechanisms.

## Acknowledgements

We thank Prof. Jun Yu (Umeå University) for helpful discussions and bioRxiv (Cold Spring Harbor Laboratory, NY, USA) for publishing preprints of the present paper (https://doi.org/10.1101/2021.07.30.454461).

## Author contributions

T.W. conceived the study, led the research, expanded FvCB models, and analysed the data. A.A., and J.S. prepared samples, and acquired gas exchange data. T.W. wrote the paper with input from all authors. T.W. revised the paper.

## Data availability statement

Data available on request from the authors.

## References

Ainsworth EA, Long SP. 2005. What have we learned from 15 years of free-air CO_2_ enrichment (FACE)? A meta-analytic review of the responses of photosynthesis, canopy properties and plant production to rising CO_2_. New Phytologist 165: 351–372.

Ayub G, Smith RA, Tissue DT, Atkin OK. 2011. Impacts of drought on leaf respiration in darkness and light in Eucalyptus saligna exposed to industrial-age atmospheric CO_2_ and growth temperature. New Phytologist 190: 1003–1018.

Bloom AJ. 2015. Photorespiration and nitrate assimilation: A major intersection between plant carbon and nitrogen. Photosynthesis Research 123: 117–128.

Buchanan BB, Gruissem W, Jones RL (Eds.). 2015. Biochemistry and molecular biology of plants. Chichester: John Wiley & Sons, Ltd.

Busch FA, Sage RF, Farquhar GD. 2018. Plants increase CO_2_ uptake by assimilating nitrogen via the photorespiratory pathway. Nature Plants 4: 46–54.

von Caemmerer S. 2000. Biochemical models of leaf photosynthesis. Collingwood: CSIRO.

Cossar JD, Rowell P, Stewart WDP. 1984. Thioredoxin as a modulator of glucose-6-phosphate dehydrogenase in a N2-fixing cyanobacterium. Microbiology 130: 991–998.

Dietz K-J. 1985. A possible rate-limiting function of chloroplast hexosemonophosphate isomerase in starch synthesis of leaves. Biochimica et Biophysica Acta 839: 240–248.

Ehlers I, Augusti A, Betson TR, Nilsson MB, Marshall JD, Schleucher J. 2015. Detecting long-term metabolic shifts using isotopomers: CO_2_-driven suppression of photorespiration in C_3_ plants over the 20^th^ century. Proceedings of the National Academy of Sciences of the United States of America 112: 15585–15590.

Eicks M, Maurino V, Knappe S, Flügge U-I, Fischer K. 2002. The plastidic pentose phosphate translocator represents a link between the cytosolic and the plastidic pentose phosphate pathways in plants. Plant Physiology 128: 512–522.

Evans J. 1987. The dependence of quantum yield on wavelength and growth irradiance. Functional Plant Biology 14: 69–79.

Evans JR. 1989. Photosynthesis and nitrogen relationships in leaves of C_3_ plants. Oecologia 78: 9–19.

Evans JR, Farquhar GD, Sharkey TD, Berry JA. 1986. Carbon isotope discrimination measured concurrently with gas exchange to investigate CO_2_ diffusion in leaves of higher plants. Australian Journal of Plant Physiology 13: 281–292.

Farquhar GD, Caemmerer S, Berry JA. 1980. A biochemical model of photosynthetic CO_2_ assimilation in leaves of C_3_species. Planta 149: 78–90.

Farquhar GD, von Caemmerer S. 1982. Modelling of photosynthetic response to environmental conditions. In: Lange OL, Nobel PS, Osmond CB, Ziegler H, eds. Physiological plant ecology II: Water relations and carbon assimilation. Berlin, Heidelberg: Springer, 549–587.

Farquhar G, Wong S. 1984. An empirical model of stomatal conductance. Functional Plant Biology 11: 191–210.

Flexas J, Diaz-Espejo A, GalméS J, Kaldenhoff R, Medrano H, Ribas-Carbo M. 2007. Rapid variations of mesophyll conductance in response to changes in CO_2_concentration around leaves. Plant, Cell & Environment 30: 1284–1298.

Genkov T, Meyer M, Griffiths H, Spreitzer RJ. 2010. Functional hybrid Rubisco enzymes with plant small subunits and algal large subunits. Journal of Biological Chemistry 285: 19833–19841.

Gerhardt R, Stitt M, Heldt HW. 1987. Subcellular metabolite levels in spinach leaves: Regulation of sucrose synthesis during diurnal alterations in photosynthetic partitioning. Plant Physiology 83: 399–407.

Gu L, Sun Y. 2014. Artefactual responses of mesophyll conductance to CO_2_and irradiance estimated with the variable J and online isotope discrimination methods. Plant, Cell & Environment 37: 1231–1249.

Harley PC, Loreto F, Di Marco G, Sharkey TD. 1992. Theoretical considerations when estimating the mesophyll conductance to CO_2_flux by analysis of the response of photosynthesis to CO_2_. Plant Physiology 98: 1429–1436.

Harley PC, Sharkey TD. 1991. An improved model of C_3_ photosynthesis at high CO_2_: Reversed O_2_ sensitivity explained by lack of glycerate reentry into the chloroplast. Photosynthesis Research 27: 169–178.

Hassiotou F, Ludwig M, Renton M, Veneklaas EJ, Evans JR. 2009. Influence of leaf dry mass per area, CO_2_, and irradiance on mesophyll conductance in sclerophylls. Journal of Experimental Botany 60: 2303–2314.

Hauschild R, von Schaewen A. 2003. Differential regulation of glucose-6-phosphate dehydrogenase isoenzyme activities in potato. Plant Physiology 133: 47–62.

Jacob J, Lawlor DW. 1991. Stomatal and mesophyll limitations of photosynthesis in phosphate deficient sunflower, maize and wheat plants. Journal of Experimental Botany 42: 1003–1011.

Kaiser E, Kromdijk J, Harbinson J, Heuvelink E, Marcelis LFM. 2017. Photosynthetic induction and its diffusional, carboxylation and electron transport processes as affected by CO_2_ partial pressure, temperature, air humidity and blue irradiance. Annals of Botany 119: 191–205.

Kruckeberg AL, Neuhaus HE, Feil R, Gottlieb LD, Stitt M. 1989. Decreased-activity mutants of phosphoglucose isomerase in the cytosol and chloroplast of Clarkia xantiana. The Biochemical Journal 261: 457–467.

Laisk A. 1977. Kinetics of photosynthesis and photorespiration in C_3_plants. Nauka, Moscow, Russia.

Laisk A, Edwards GE. 1997. CO2 and temperature-dependent induction in C_4_ photosynthesis: an approach to the hierarchy of rate-limiting processes. Functional Plant Biology 24: 505–516.

Liu T, Barbour MM, Yu D, Rao S, Song X. 2022. Mesophyll conductance exerts a significant limitation on photosynthesis during light induction. New Phytologist 233: 360–372.

Nadal M, Carriquí M, Flexas J. 2021. Chapter 3 Mesophyll conductance to CO_2_ diffusion in a climate change scenario: Effects of elevated CO_2_, temperature and water stress. In: Becklin KM, Ward JK, Way DA, eds. Photosynthesis, Respiration, and Climate Change. Cham: Springer International Publishing, 49–78.

Orr DJ, Alcântara A, Kapralov MV, Andralojc PJ, Carmo-Silva E, Parry MAJ. 2016. Surveying Rubisco diversity and temperature response to improve crop photosynthetic efficiency. Plant Physiology 172: 707–717.

Preiser AL, Fisher N, Banerjee A, Sharkey TD. 2019. Plastidic glucose-6-phosphate dehydrogenases are regulated to maintain activity in the light. Biochemical Journal 476: 1539–1551.

Sakoda K, Yamori W, Groszmann M, Evans JR. 2021. Stomatal, mesophyll conductance, and biochemical limitations to photosynthesis during induction. Plant Physiology 185: 146–160.

Schäufele R, Santrucek J, Schnyder H. 2011. Dynamic changes of canopy-scale mesophyll conductance to CO_2_ diffusion of sunflower as affected by CO_2_ concentration and abscisic acid. Plant, Cell & Environment 34: 127–136.

Scheibe R. 1981. Thioredoxinm in pea chloroplasts: Concentration and redox state under light and dark conditions. FEBS Letters 133: 301–304.

Scheibe R. 2004. Malate valves to balance cellular energy supply. Physiologia Plantarum 120: 21–26.

Schleucher J, Vanderveer P, Markley JL, Sharkey TD. 1999. Intramolecular deuterium distributions reveal disequilibrium of chloroplast phosphoglucose isomerase. Plant, Cell and Environment 22: 525–533.

Seelert H, Poetsch A, Dencher NA, Engel A, Stahlberg H, Müller DJ. 2000. Proton-powered turbine of a plant motor. Nature 405: 418–419.

Sharkey TD. 1985. O_2_-insensitive photosynthesis in C_3_ plants: Its occurrence and a possible explanation. Plant Physiology 78: 71–75.

Sharkey TD, Berry JA, Raschke K. 1985. Starch and sucrose synthesis in Phaseolus vulgaris as affected by light, CO_2_, and abscisic acid. Plant Physiology 77: 617–620.

Sharkey TD, Weise SE. 2016. The glucose 6-phosphate shunt around the Calvin–Benson cycle. Journal of Experimental Botany 67: 4067–4077.

Steigmiller S, Turina P, Gräber P. 2008. The thermodynamic H+/ATP ratios of the H+-ATPsynthases from chloroplasts and Escherichia coli. Proceedings of the National Academy of Sciences of the United States of America 105: 3745–3750.

Taira M, Valtersson U, Burkhardt B, Ludwig RA. 2004. Arabidopsis thaliana GLN2-encoded glutamine synthetase is dual targeted to leaf mitochondria and chloroplasts. The Plant Cell 16: 2048–2058.

Tcherkez G, Gauthier P, Buckley TN, Busch FA, Barbour MM, Bruhn D, Heskel MA, Gong XY, Crous KY, Griffin K, et al. 2017. Leaf day respiration: low CO_2_ flux but high significance for metabolism and carbon balance. New Phytologist 216: 986–1001.

Tcherkez G, Limami AM. 2019. Net photosynthetic CO_2_ assimilation: more than just CO_2_ and O_2_ reduction cycles. New Phytologist 223: 520–529.

Tholen D, Ethier G, Genty B, Pepin S, Zhu X-G. 2012. Variable mesophyll conductance revisited: theoretical background and experimental implications. Plant, Cell & Environment 35: 2087–2103.

Tjoelker MG, Oleksyn J, Lorenc-Plucinska G, Reich PB. 2009. Acclimation of respiratory temperature responses in northern and southern populations of Pinus banksiana. New Phytologist 181: 218–229.

Vrábl D, Vašková M, Hronková M, Flexas J, Šantrůček J. 2009. Mesophyll conductance to CO_2_ transport estimated by two independent methods: effect of variable CO_2_ concentration and abscisic acid. Journal of Experimental Botany 60: 2315–2323.

Walker AP, Beckerman AP, Gu L, Kattge J, Cernusak LA, Domingues TF, Scales JC, Wohlfahrt G, Wullschleger SD, Woodward FI. 2014. The relationship of leaf photosynthetic traits - Vcmax and Jmax - to leaf nitrogen, leaf phosphorus, and specific leaf area: a meta-analysis and modeling study. Ecology and evolution 4: 3218–3235.

Wang X, Lewis JD, Tissue DT, Seemann JR, Griffin KL. 2001. Effects of elevated atmospheric CO_2_ concentration on leaf dark respiration of Xanthium strumarium in light and in darkness. Proceedings of the National Academy of Sciences of the United States of America 98: 2479–2484.

Weise SE, Schrader SM, Kleinbeck KR, Sharkey TD. 2006. Carbon balance and circadian regulation of hydrolytic and phosphorolytic breakdown of transitory starch. Plant Physiology 141: 879–886.

Wieloch T. 2021. The next phase in the development of 13C isotopically nonstationary metabolic flux analysis. Journal of Experimental Botany 72: 6087–6090.

Wieloch T. 2022. High atmospheric CO2 concentration causes increased respiration by the oxidative pentose phosphate pathway in chloroplasts. New Phytologist 235: 1310–1314.

Wieloch T, Augusti A, Schleucher J. 2022a. Anaplerotic flux into the Calvin-Benson cycle. Hydrogen isotope evidence for in vivo occurrence in C_3_ metabolism. New Phytologist 234: 405–411.

Wieloch T, Ehlers I, Yu J, Frank D, Grabner M, Gessler A, Schleucher J. 2018. Intramolecular 13C analysis of tree rings provides multiple plant ecophysiology signals covering decades. Scientific Reports 8: 5048.

Wieloch T, Grabner M, Augusti A, Serk H, Ehlers I, Yu J, Schleucher J. 2022b. Metabolism is a major driver of hydrogen isotope fractionation recorded in tree-ring glucose of Pinus nigra. New Phytologist 234: 449–461.

Wieloch T, Sharkey TD. 2022. Compartment-specific energy requirements of photosynthetic carbon metabolism in Camelina sativa leaves. Planta 255: 103.

Wullschleger SD. 1993. Biochemical limitations to carbon assimilation in C3 plants. A retrospective analysis of the A/Ci curves from 109 Species. Journal of Experimental Botany 44: 907–920.

Xiong D, Liu X, Liu L, Douthe C, Li Y, Peng S, Huang J. 2015. Rapid responses of mesophyll conductance to changes of CO2 concentration, temperature and irradiance are affected by N supplements in rice. Plant, Cell & Environment 38: 2541–2550.

Xu Y, Fu X, Sharkey TD, Shachar-Hill Y, Walker BJ. 2021. The metabolic origins of non-photorespiratory CO2 release during photosynthesis: A metabolic flux analysis. Plant Physiology 186: 297–314.

Xu Y, Wieloch T, Kaste JAM, Shachar-Hill Y, Sharkey TD. 2022. Reimport of carbon from cytosolic and vacuolar sugar pools into the Calvin–Benson cycle explains photosynthesis labeling anomalies. Proceedings of the National Academy of Sciences of the United States of America 119: e2121531119.

Yin X, van der Putten PEL, Belay D, Struik PC. 2020. Using photorespiratory oxygen response to analyse leaf mesophyll resistance. Photosynthesis Research 144: 85–99.

Zeeman SC, Thorneycroft D, Schupp N, Chapple A, Weck M, Dunstan H, Haldimann P, Bechtold N, Smith AM, Smith SM. 2004. Plastidial α-glucan phosphorylase is not required for starch degradation in Arabidopsis leaves but has a role in the tolerance of abiotic stress. Plant Physiology 135: 849–858.

