## Supporting information for "A model of photosynthetic CO_2_ assimilation in C_3_ leaves accounting for respiration and energy recycling by the plastidial oxidative pentose phosphate pathway"

**Notes S1. Discussion of model parameter estimates including sensitivity analyses**

Modelling results pertaining to RuBP-regeneration-limited $CO_2$ assimilation rely on the parameterisation of $\Gamma_*$ and $J_{max}$ (Eqs. 12, and 25). In $C_3$ plants at 21% $O_2$, $\Gamma_*$ is usually within 35 to 45 µbar $CO_2$. The value we calculated is at the upper limit ($\Gamma_*$ = 45.3 µbar $CO_2$). Changing $\Gamma_*$ in our $A_j$ model to 55.2 µbar $CO_2$ (while keeping everything else unchanged) results in negligible offsets between $A_j$ and $A_m$ and thus negligible $R_{a,p}$ at low $C_a$ (Fig. S1). However, this value is unusually high, and the resulting $A_j$ model does not capture $A_m$ values measured at 450 and 700 ppm well. Therefore, our modelling results ($R_{a,p}$) are qualitatively robust with respect to our parameterisation of $\Gamma_*$.

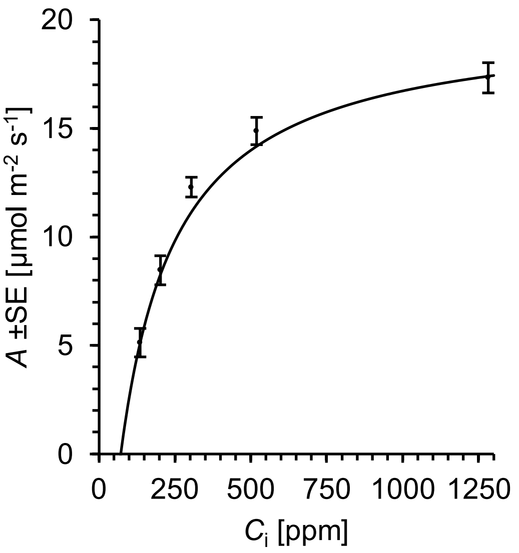


**Figure S1** Net $CO_2$ assimilation ($A$) of sunflower leaves as function of intercellular $CO_2$ concentration ($C_i$). Black dots: measured values ($A_m$, $n$ = 8). Solid black line: modelled RuBP-regeneration-limited assimilation ($A_j$) (respiration by the plastidial anaplerotic pathway not considered). Plants were grown in a greenhouse at an atmospheric $CO_2$ concentration ($C_a$) of ≈ 450 ppm and then moved to growth chambers. After a day in darkness to drain the starch reserves, plants were grown at different levels of $C_a$ (180, 280, 450, 700, 1500 ppm) corresponding to different levels of $C_i$ (140, 206, 328, 531, and 1365 ppm) for two days. During gas exchange measurements slightly lower $C_i$ levels prevailed (136, 202, 304, 516, 1282 ppm). $A_j$-model inputs: $\Gamma_*$ = 55.2 µbar $CO_2$, $J_{max}$ = 168.3 µmol $m^{-2}$ $s^{-1}$, θ = 0.7, $I$ = 300 µmol photons $m^{-2}$ $s^{-1}$, abs = 0.85, and f = 0.15.

In the present study, $J_{max}$ was estimated from $V_{cmax}$ based on a reported statistical relationship between these parameters (Eqn. 27). In turn, $V_{cmax}$ was estimated using reported rubisco kinetic parameters and $g_m$ (Fig. 3). Hence, our $J_{max}$ estimate can be expected to suffer from (i) measurement/estimation error of rubisco kinetic parameters and $g_m$, (ii) error due to parameter differences among sunflower cultivars and environmental conditions, (iii) error due to the temperature correction of parameter estimates, (iv) error propagation on estimating $V_{cmax}$, and (v) error related to the statistical relationship between $V_{cmax}$ and $J_{max}$. That said, we estimated $J_{max}$ = 168.3 µmol $m^{-2}$ $s^{-1}$. To assess how sensitive our results are to changes in $J_{max}$, we recalculated main results after adjusting $J_{max}$ at 150, 160, 170, 180, and 190 µmol $m^{-2}$ $s^{-1}$. We found that our estimates of $R_{a,p}$ and $R_x$ are qualitatively robust over this $J_{max}$ range (Table S1). However, with decreasing $J_{max}$ below 150 µmol $m^{-2}$ $s^{-1}$, $R_x$ moves towards and below zero.

**Table S1** Sensitivity of $R_{a,p}$ and $R_x$ estimates to changes in $J_{max}$.

| $J_{max}$ [µmol $m^{-2}$ $s^{-1}$] | $R_{a,p}$ [µmol $m^{-2}$ $s^{-1}$] | | $R_x$ [µmol $m^{-2}$ $s^{-1}$] | | |
|---|---|---|---|---|---|
| | 180 ppm | 280 ppm | 450 ppm | 700 ppm | 1500 ppm |
| 168.3 | 2.75 | 2.44 | 0.96 | 1.08 | 1.38 |
| 150 | 3.08 | 2.64 | 0.46 | 0.47 | 0.67 |
| 160 | 2.90 | 2.53 | 0.75 | 0.82 | 1.08 |
| 170 | 2.73 | 2.43 | 1.00 | 1.13 | 1.44 |
| 180 | 2.58 | 2.33 | 1.23 | 1.40 | 1.76 |
| 190 | 2.44 | 2.25 | 1.44 | 1.65 | 2.05 |

Modelling results pertaining to rubisco-limited $CO_2$ assimilation rely on the parameterisation of $K_c$, $K_o$, $S_{c/o}$, and $V_{cmax}$ (Eqs. 5, and 24). These estimates vary strongly among species (Orr *et al.*, 2016). Here, we used values for sunflower rubisco reported by Genkov *et al.* (2010). To our knowledge, this is the only complete sunflower datasets. That is, there is a lack of knowledge about parameter variability among sunflower cultivars and environmental conditions. To complicate matters further, rubisco kinetic parameters are partially intercorrelated (Orr *et al.*, 2016), and can be expected to exert interactive effects on $A_c$ (Eqn. 5). That is, sensitivity analysis with respect to these parameters is not straightforward since varying parameters independently would be too simplistic. Therefore, tests for robustness of our results with respect to rubisco kinetic parameters are practically unfeasible. Considering uncertainties related to rubisco kinetic parameters, results reported here require validation. Hence, we recommend follow-up studies that combine gas exchange measurements, isotope measurements, and model parameter measurements from the same plants.

**References**

**Genkov T, Meyer M, Griffiths H, Spreitzer RJ**. **2010**. Functional hybrid Rubisco enzymes with plant small subunits and algal large subunits. *Journal of Biological Chemistry* **285**: 19833–19841.

**Orr DJ, Alcântara A, Kapralov MV, Andralojc PJ, Carmo-Silva E, Parry MAJ**. **2016**. Surveying Rubisco diversity and temperature response to improve crop photosynthetic efficiency. *Plant Physiology* **172**: 707–717.
